# Adaptable centriole biogenesis via the intrinsically disordered protein ALMS1

**DOI:** 10.1101/2025.06.26.661604

**Authors:** Kanako Ozaki, Ting-Jui Ben Chang, Wen-Qing Yang, Avital Shulman, Denisse Izquierdo, Wann-Neng Jane, Won-Jing Wang, Tim Stearns, Jens Lüders, T. Tony Yang, Meng-Fu Bryan Tsou

**Affiliations:** Cell Biology Program, Sloan Kettering Institute, Memorial Sloan Kettering Cancer Center, New York; Department of Electrical Engineering, National Taiwan University, Taipei, Taiwan; Weill Cornell Graduate School of Medical Sciences, Cornell University, New York; Plant Cell Biology Center, Institute of Plant and Microbiology, Sinica Academia, Taipei, Taiwan; Institute of Biochemistry, National Yang Ming Chiao Tung University, Taipei, Taiwan; Head of the Laboratory of Cellular Dynamics, The Rockefeller University, New York; Institute for Research in Biomedicine (IRB Barcelona), The Barcelona Institute of Science and Technology, Barcelona, Spain; Graduate Institute of Biomedical Electronics and Bioinformatics, National Taiwan University, Taiwan

## Abstract

Biogenesis of subcellular structures like centrioles is viewed as a physical transformation wherein elementary constituents form order without preexisting templates. Centrioles grow with precision from a composite scaffold known as the cartwheel, which is thought to self-assemble without templates and disassemble following centriole growth; however, the mechanism governing cartwheel assembly-disassembly dynamics remains obscure. Here, we identify ALMS1, a disease-linked, intrinsically disordered protein (IDP), as an external mediator of cartwheel dynamics that causes a seed for cartwheel—and thus centriole—formation without itself incorporating into the seed structure. The cartwheel seed (CS), characterized as a dense composite of CEP152/CEP63 protein complexes, forms in interphase and adopts a nanoscale, concentric ring from which the cartwheel grows. Upon mitotic entry, CSs recruit ALMS1 while disassembling into constituents associating with ALMS1 in proximity, correlating with cartwheel assembly-disassembly cycles. Hypomorph, disease-linked ALMS1 mutations trigger cartwheel expansion and shedding by its own grown procentriole, in turn forming ectopic centrioles, leading to perpetual reciprocal amplification. Without ALMS1, CS formation fails, negating centriole biogenesis, whereas reintroducing ALMS1 initializes biogenesis anew, creating diverse yet heritable architectures that evolve through selection, instead of generating a single canonical form. These results suggest that centriole biogenesis is grounded on adaptable transformation cues extrinsic to its constituents, propagating via IDP-mediated CS assembly-disassembly cycles, a process we conjecture involves memory.

## Introduction

The emergence of composite objects from elementary constituents in biogenesis is a poorly understood physical transformation in which an initial, orderless configuration of substrates and energy is mapped onto a distinct, functional output configuration. Transformation cues that cause biogenesis may be solely intrinsic to physicochemical properties of the constituents or, alternatively, imposed by extrinsic factors that do not themselves incorporate into the final composite object. In protein synthesis, for instance, transforming amino acids into specific polypeptides is directed by external nucleic acid templates that are themselves adaptive and transmittable. Such external information systems allow turnovers of the constituents, composite products, or both without losing the cues for resuming identical transformations when needed.

In contrast, at the subcellular level, the assembly of composite objects like centriole organelles from protein-based constituents is generally thought to occur de novo, without external templates, via spontaneous interactions among constituents. This widely accepted view, while lacking critical empirical support, regards centrioles exclusively as a genetically determined product capable of emerging with high precision and negligible plasticity. It is known that centrioles in some animals can be naturally eliminated and then regenerated as needed in the absence of preexisting copies^1–5^, like protein turnover. However, whether centriole assembly also involves external cues or solely depends on its constituents remains an open question. Here, we show that centriole biogenesis is caused by adaptable transformation cues external to its constituents, capable of creating diverse yet heritable architectures.

Centrioles grow with an invariable shape from the cartwheel scaffold, a stack of ring-like structures each comprising, among other proteins, SAS-6 molecules arranged in 9-fold radial symmetry^6–9^. The cartwheel appears to form de novo in interphase, either at multiple locations when cells lack preexisting centrioles or, alternatively, at one site around each existing (or mother) centriole. In both cases, each cartwheel grows into a new (or daughter) centriole. The SAS-6 in the cartwheel turns over much faster than the centriole backbone^10–12^, implying that in human cells the cartwheel scaffold is a dynamic rather than an integral part of the organelle. PLK4 and its substrate STIL activate the assembly^13^, with both incorporating into the cartwheel, essential for stability^14^. PLK4, which activates cartwheel formation in canonical duplication, is recruited by centriole-associated CEP152/CEP63 complexes^15–18^. Newborn daughter centrioles cannot re-duplicate until going through centriole-to-centrosome conversion (CCC) in late G2 and mitosis (G2/M)^19–21^. Through CCC, newborn centrioles acquire the pericentriolar material (PCM) containing CEP152/CEP63 complexes, enabling the duplication potential. Notably, coinciding with CCC, the cartwheel embedded in newborn centrioles disassembles, involving mitotic kinases PLK1^19^ and CDK1^12, 22^, generating mature centrioles devoid of the cartwheel to start G1. What underlies cartwheel dynamics remains unclear, leading to a prevailing view that cartwheels self-assemble from orderless constituents without preexisting templates. Here, we show that the cartwheel is instead grown by a seed that forms via an intrinsically disordered mediator—the Alström syndrome protein ALMS1^23, 24^—which is not part of the seed or cartwheel structure.

## Results

### Dysregulation of cartwheel dynamics in G2/M causes premature cartwheel shedding and abnormal centriole growth

To characterize cartwheel dynamics during the cell cycle, cells arrested in S or G2 and cells released into M from G2 arrest were examined with ultrastructure-optimized expansion microscopy (UExM)^25^. We found that the cartwheel in the centriole lumen grows about twofold in length from S- to G2-arrested conditions (Fig. 1a) and that upon mitotic entry from the G2 arrest, the cartwheel signal diminishes quickly in a PLK1- but not proteosome-dependent manner (Fig. 1b), becoming nearly undetectable by UExM at metaphase (Fig. 1b). CDK1, the kinase that drives mitotic entry, is known to regulate cartwheel removal^22^. We reasoned that perhaps reducing CDK1 activities to slow down the G2-to-M progression may reveal new details of cartwheel dynamics before the cartwheel disappears. To test the idea, we developed cell synchronization assays for quantitative studies based on a published protocol^26^. In brief, *NF2^-/-^; p53^-/-^* RPE1 cells, which lack proper contact inhibition and stress response, can grow synchronously from G1 to G2 at high cell densities (see Materials and Methods; Fig. S1a), a condition essential for our UExM studies. Treating cells synchronously entering G2 with strong CDK1 inhibition (9.5 μM RO-3306) fully blocked mitotic entry for a 12-hour time course of live cell imaging (Fig. S1a); conversely, a treatment with partial CDK1 inhibition (6 μM RO-3306) allowed cells to slowly progress into mitosis (Fig. S1a). Under this 12-hour time course during which cells slowly progress toward and over the G2/M transition, cartwheel dynamics were analyzed by UExM. We found that over time, but before cells enter mitosis, the cartwheel gradually transforms from being intact in the lumen (INT) to a dumbbell-like shape (DBL), as if the cartwheel were dividing, then to what we termed the split-and-shed configuration (SHD)—where the cartwheel appears to split into two separate units, with one shedding from the lumen (Fig. 1c). The percentage of intact cartwheels drops from 79.6% at 3hr to 30.4% at the 12h timepoint, whereas cartwheel shedding increases from 0% at 3hr to 32.7%r at 12h. No cartwheel shedding was seen in control cells fully arrested in G2 by strong CDK1 inhibition (Fig. 1c).

**Fig. 1.**
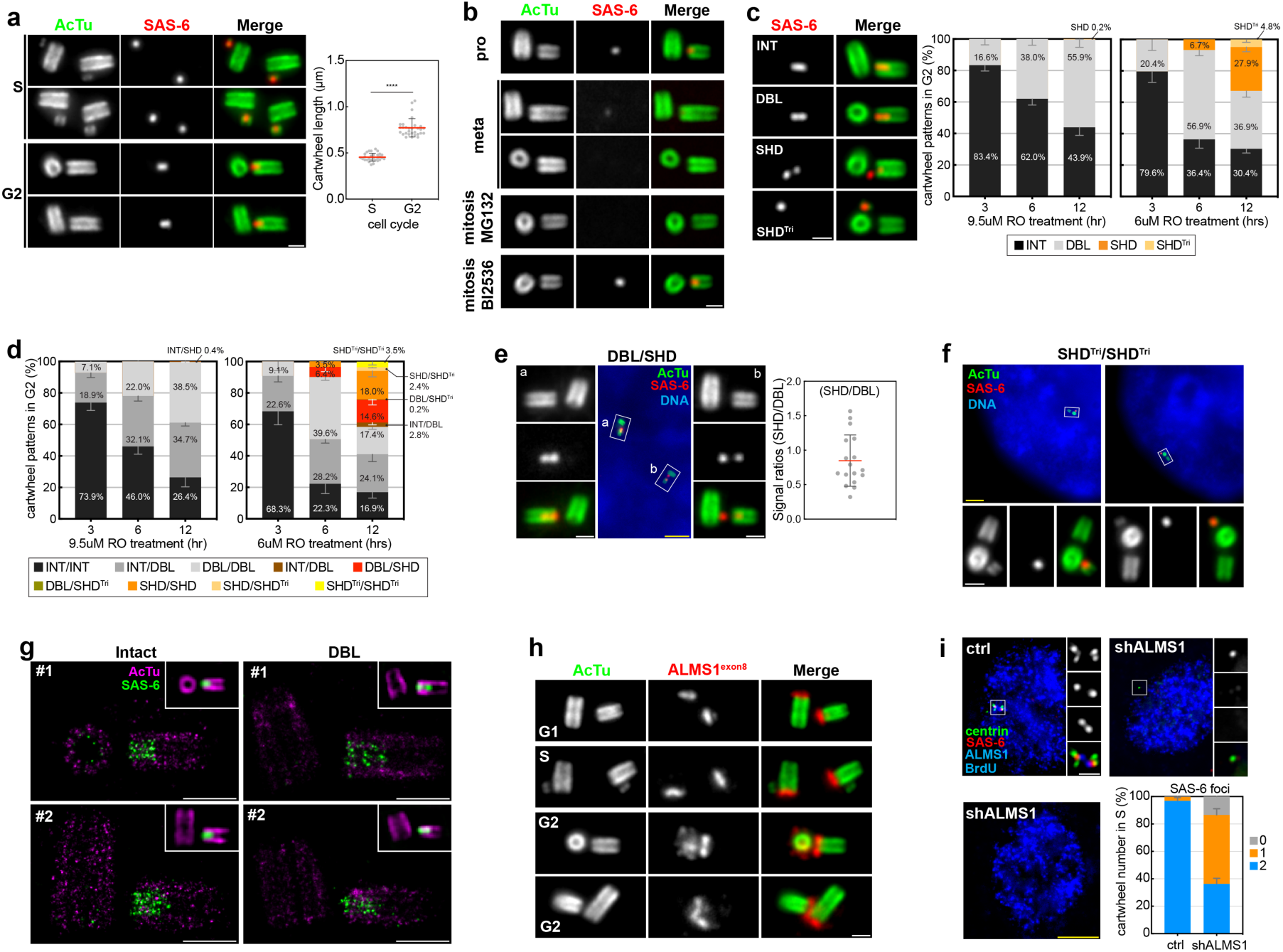
Dysregulation of G2/M cartwheel dynamics causing abnormal centriole growth, and identification of ALMS1 as a non-constituent of the cartwheel essential for cartwheel formation. (**a**) UExM analyses of the centriole (acetylated α-tubulin/AcTu; green) and cartwheel (SAS-6; red) arrested at S or G2 phase. S- or G2-phase centrioles were examined in RPE1 cells treated with aphidicolin (2 μg/ml) or RO-3306 (9.5 μM) for 20 hrs. Quantification of the cartwheel length was done with samples from three independent experiments. Data are mean ± s.d., analyzed by Welch’s t-test (p<0.01). (**b**) UExM images of the centriole and cartwheel at indicated phases of mitosis with or without indicated drug treatments. Mitotic cells were obtained by releasing G2-arrested cells into the cell cycle for 45min. The proteasome inhibitor MG132 (10 μM) or PLK1 inhibitor BI-2536 (200 nM) was used. (**c**) Representative images of cartwheel transformation from being intact (INT) to dumbbell-like (DBL), split-and-shed (SHD), and shed with centriole triplets (SHD^Tri^) in G2 cells growing in the presence of strong (9.5 μM RO) or mild (6 μM RO) CDK1 inhibition (bottom). Quantitation was shown. Data are mean ± s.d., with N = 3 independent biological replicates. (**d**) Reanalyzing cartwheel shedding in (c) by grouping the two pairs of centrioles from the same cell as a data point. Cartwheel shedding in each combination was indicated. Data are mean ± s.d., with N = 3 independent biological replicates. (**e**) UExM images and quantification of the two cartwheel structures in the same cell, with one exhibiting SHD and the other DBL configurations (SHD/DBL). Signal ratios were obtained by dividing the total signal of SHD cartwheels (two split units) with that of DBL cartwheels. No net increase in cartwheel signal was seen upon shedding. (**f**) UExM images of the two pairs of centriole triplets in the same cell under partial CDK1 inhibition. The shed cartwheel that tethers to mother centrioles appeared to grow into procentrioles, generating abnormal centriole triplet configurations. (**g**) Ex-dSTORM analyses of the cartwheel (SAS-6; green) before and after it changes to DBL in the lumen. Two intact and DBL luminal cartwheels (#1-#2) are shown. Scale bars, 1 μm (white). (**h**) UExM analyses of ALMS1 localization using antibodies against epitopes encoded by ALMS1 exon 8. Antibodies against exon1, exon14/15, or exon23 were shown in Fig. S1. ALMS1 was not seen in S-phase procentrioles but localized to the proximal end of G2 procentrioles. (**i**) RPE1 cells treated with shRNA for 84 hrs were examined for ALMS1 localization and centriole duplication in S phase. Depletion of ALMS1 was confirmed. Failure of cartwheel and procentriole formation in *shALMS1* cells in S phase (BrdU) was seen, as revealed by centrin and SAS-6 staining. Quantification is shown. Data are mean ± s.d., with N = 3 independent biological replicates. All DNA was shown in blue (DAPI). All scale bars are 1 μm (white) and 5 μm (yellow).

G2 cells normally host two cartwheel structures in two separate newborn daughters. Analyzing the two cartwheels together as a pair in individual cells through the time course revealed progressive transformations, from both cartwheels being intact (INT/INT) to INT/DBL, DBL/DBL, DBL/SHD, then SHD/SHD (Fig. 1d; Fig. S1b), indicating that while the two cartwheels in the same cell progress closely through the same transformation, they are not completely in sync. We then analyzed individual DBL/SHD cells in which one cartwheel had split and shed while the other remained in the lumen or unshed. We found that there are no consistent net increases in SAS-6 signals between the DBL and SHD cartwheel (Fig. 1e), supporting the idea that the transformation of DBL into SHD is not a result of additional cartwheel growth. Intriguingly, among the shed cartwheels, which are tethered to the engaged mother centriole, ∼13% of them appeared to grow into a procentriole (Fig. 1c & d, SHD^Tri^), producing a striking centriole triplet in which the mother centriole is simultaneously engaged to a full-length daughter devoid of the cartwheel and a short procentriole carrying the cartwheel (Fig. 1f; Fig. S1b). Interestingly, similar centriole triplets were identified in *Drosophila* tissues exhibiting diminished CDK1 activities^27^, suggesting the specificity of this phenotype caused by partial CDK1 inhibition.

To reveal the detailed organization of the cartwheel before shedding, the side view of the cartwheel in the lumen before and after it changes to DBL shapes under UExM was correlatively examined by direct stochastic optical reconstruction microscopy, or Ex-dSTORM^28^. We found that intact cartwheels identified under UExM also show uniform SAS-6 staining along the length of the structure under dSTORM (Fig. 1g, Intact). In contrast, Ex-dSTORM analyses of DBL cartwheels revealed the presence of a gap in between two parts that appear to be spatially disoriented from each other (Fig. 1g; DBL), supporting the idea that under partial CDK1 inhibition the expanded cartwheel can physically split in the lumen before shedding.

Furthermore, examining cells at the 12hr timepoint that had crossed the G2/M border and were progressing in mitosis under partial CDK1 inhibition revealed that cartwheel shedding or disassembly occurs in >95% of mitotic cells (Fig. S1c), with the cartwheel number count ranging from 2 to 0 per mother-daughter centriole pair (Fig. S1c & S1d). A representative mitotic cell showing two shed cartwheels per mother-daughter centriole pair, the (2, 2) configuration, was shown (Fig. S1d). By G1, all cartwheel structures were lost (Fig. S1e), suggesting that cartwheel disassembly, despite being slower, still occurs under partial CDK1 inhibition. Together, these results suggest that cartwheel dynamics during G2/M are under regulation and that their dysregulation by partial CDK1 inhibition can lead to abnormal centriole growth before mitotic entry.

### Identification of ALMS1 as a non-constituent of the cartwheel essential for cartwheel formation

To identify novel regulators of cartwheel dynamics, we searched for molecules that are absent at the procentriole or cartwheel during early S but present there during G2. UExM analyses of the Alström syndrome protein ALMS1^23^ using antibodies against 4 different regions revealed that ALMS1 is undetectable at S-phase procentrioles but becomes highly enriched at the proximal end of procentrioles during G2 (Fig. 1h; Fig. S1f). Examining ALMS1 in free-standing procentrioles that form in the absence of preexisting centrioles in *p53^-/-^; CEP295^-/-^* cells^20^ showed a similar pattern, with ALMS1 being undetectable until G2 (Fig. S1g). Knocking down ALMS1 in p53-null cells known to tolerate centrosome loss^29^ using siRNA (Fig. S1h) or shRNA (Fig. 1i) for 3-4 days revealed that cartwheel or procentriole assembly is blocked in >60% of cells in S phase (BrdU), with some of these S-phase cells completely devoid of cartwheel- or procentriole-like structures (Fig. 1i). The similar requirement of ALMS1 for cartwheel formation has recently been reported in *Drosophila* cells^30^. Surprisingly, long-term depletion of ALMS1 by shRNA severely disrupted spindle assembly (Fig. S1i) and stopped cell proliferation even in the absence of p53 (not shown), an intriguing phenotype that is not explainable by centrosome loss alone and will be studied elsewhere. Similar cell cycle arrest was seen with CRISPR targeting exon 1 of ALMS1, rendering isolation of ALMS1 knockout clones unfeasible. Cells freshly deleted of ALMS1 at day 5-6 after CRISPR treatments were examined for cartwheel assembly (Fig. S1j). Among cells in which ALMS1 deletion had led to complete centrosome loss, as judged by the lack of γ-tubulin foci, no cartwheel or procentriole can be detected in S phase (Fig. S1j). Thus, while ALMS1 is undetectable at the cartwheel or procentrioles in the S phase, it is required for their formation.

### Disease-associated mutations in ALMS1 induce reciprocal amplification of the cartwheel and centriole via a positive feedback loop

We next examined if ALMS1-hypomorph mutations associated with the disease have impacts on cartwheel dynamics^31, 32^. *p53^-/-^* cells introduced with homozygous nonsense mutations at exon8 of ALMS1 (*ALMS1^Δex8^*) were found to carry normal-looking centrioles capable of undergoing duplication (Fig. S2a) and recruiting ALMS1 isoforms lacking exon8 epitopes (Fig. S2b). Arresting *ALMS1^Δex8^* cells at S phase did not significantly change the cartwheel or centriole number (Fig. 2a). In contrast, inducing G2 arrest by treating asynchronous *ALMS1^Δex8^* cells with strong CDK1 inhibition (9.5 µM RO3306) for 24hrs caused cartwheel amplifications in >60% of cells (Fig. 2a). Carefully examining the extent of cartwheel amplification in G2-arrested cells revealed patterns (Fig. S2c-p). To better describe the pattern, we introduced a 1x3 matrix representation to classify cartwheel amplification in each G2-arrested *ALMS1^Δex8^* cell. This matrix quantifies the total number of cartwheels per cell (1^st^ #), of which the number of cartwheels associated with engaged centriole doublets (2^nd^ #) or with disengaged centriole singlets (3^rd^ #) was separately recorded. For example, wild-type (WT) cells arrested at G2 carry <u>two</u> cartwheels, of which <u>two</u> are associated with engaged centrioles and <u>zero</u> with disengaged centrioles, giving rise to the “2-2-0” configuration (Fig. S2c). Using this matrix, cartwheel amplification in G2-arrested *ALMS1^Δex8^* cells was orderly classified into 5 groups—as defined by the total number of cartwheels—in 12 configurations (Fig. S2c-p; summarized in Fig. 2b), with the following observations: (i) The number of cartwheels can increase ahead of centriole numbers but not vice versa (Fig. S2d, e, g, & h), (ii) every cartwheel, ranging from 2 to 6 per cell, is associated with either a disengaged centriole (singlet) or a pair of engaged centrioles (doublet) (Fig. S2c-p), and (iii) while the total number of cartwheels per cell varies, the number of active centrosomes, indicated by bright γ-tubulin signals at the PCM, remains unchanged at two per cell (Fig. S2c-p). These data suggest that in G2-arrested *ALMS1^Δex8^* cells, extra cartwheels form to seed new centrioles, but as the cell cycle is paused at late G2, newly formed centrioles are not converted to centrosomes.

**Fig. 2.**
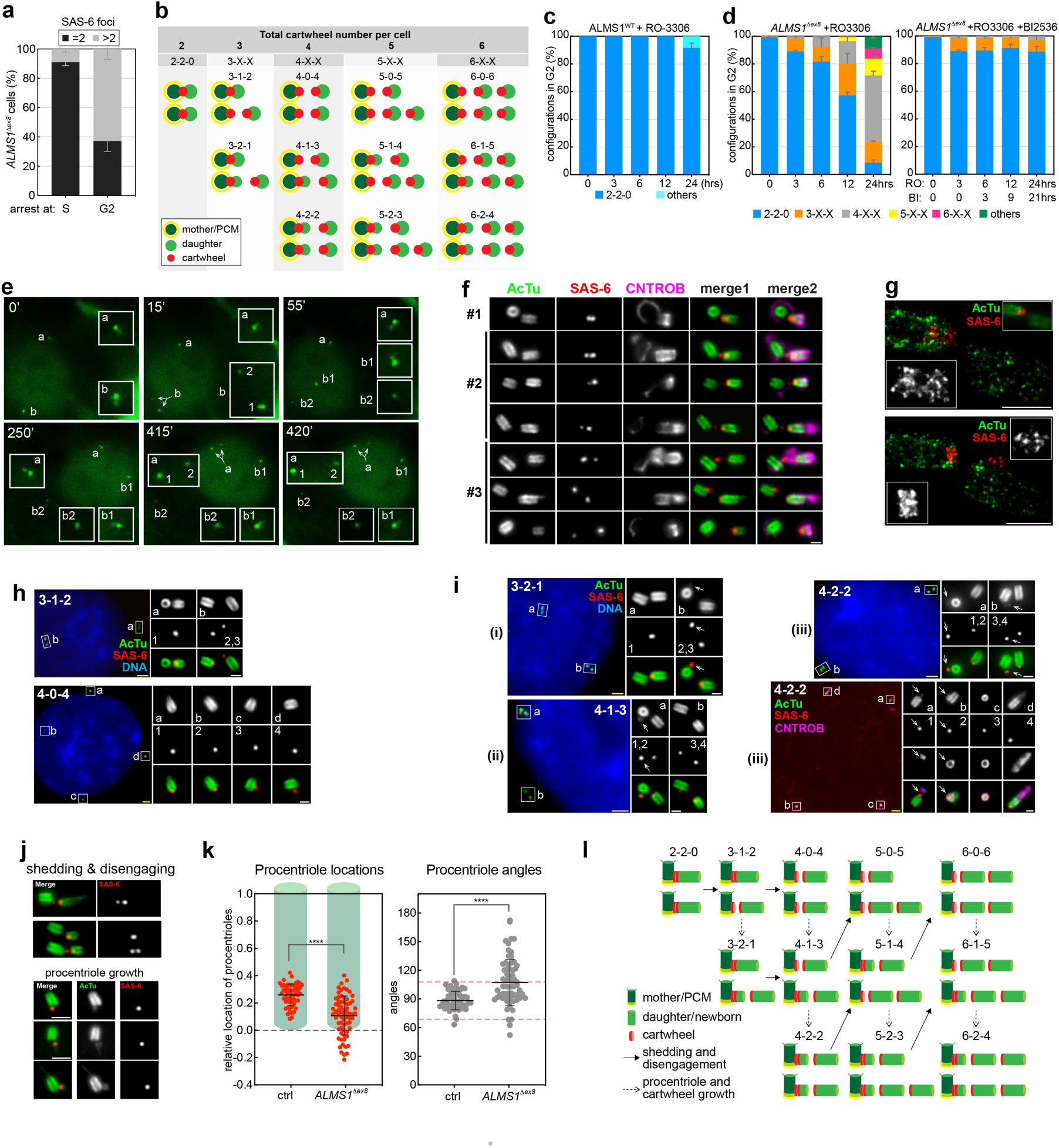
Disease-associated mutations in ALMS1 induce reciprocal amplification of the cartwheel and centriole via a positive feedback loop. (**a**) Quantification of SAS-6 foci in *ALMS1^Δex8^*cells arrested at S or G2 phase. S or G2 cells were enriched by treating unsynchronized *ALMS1^Δex8^* cells with aphidicolin (2 μg/ml) or RO-3306 (9.5 μM), respectively, for 20 hrs. Data are mean ± s.d., with N = 3 independent biological replicates. (**b**) Summary of all forms of cartwheel amplification seen in G2-arrested *ALMS1^Δex8^* cells from (**a**). The mother centriole (dark green), daughter centriole (light green), cartwheel (red), and PCM (yellow) were marked as indicated. Cartwheel amplification in each cell could be classified into one of the 12 configurations in 5 groups using 1x3 matrix representations, with the first number indicating the total number of cartwheels per cell, the second indicating the number of cartwheels associating with engaged centrioles (green doublets), and the third indicating the number of cartwheels associating with disengaged centrioles (green singlets). The phenotype was highly patterned (see the main text). (**c** and **d**) Time-course analyses of cartwheel dynamics in ALMS1^WT^ (**c**) or *ALMS1^Δex8^* (**d**) cells fully arrested in G2. Cells synchronously entering G2 were treated with RO-3306 (9.5 μM) for indicated time and fixed. No cartwheel amplification was seen in control ALMS1^WT^ cells (**c**). Cartwheel amplification that occurred over time and in specific configurations was seen in *ALMS1^Δex8^* cells (**d**, left), which was completely suppressed by PLK1 inhibition (200 nM BI-2536) applied 3 hrs after adding RO-3306 (**d**, right). Data are mean ± s.d., with N = 3 independent biological replicates. (**e**) *ALMS1^Δex8^* cells expressing centrin:GFP and synchronously entering and being arrested at G2 for 6 hrs by RO-3306 (9.5 μM) were filmed by time-lapse microscopy for an additional 10 hrs. Still images of a G2 cell carrying two pairs of engaged centrioles undergoing disengagement were extracted from the movie. Split arrows indicate centriole disengagement. Box b1 at the 55min timepoint marks reduplication. (**f**) Early phases of cartwheel splitting and shedding that accompany centriole disengagement in *NF2^-/-^; p53^-/-^; ALMS1^Δex8^* cells synchronously arrested at G2 for 6 or 12 hrs. Daughter centrioles disengaging from mothers were marked by CNTROB labeling (magenta). (**g**) Ex-dSTORM analyses of the cartwheel splitting and shedding in *ALMS1^Δex8^* cells arrested in G2. Top, a pair of *ALMS1^Δex8^*centrioles initiating cartwheel shedding, accompanying centriole disengagement. Note that the cartwheel had physically split, with the proximal unit partially exiting the lumen. Bottom, a pair of *ALMS1^Δex8^* centrioles that had shed one cartwheel unit while keeping the other unit in the lumen. (**h**) UExM images of the 3-1-2 and 4-0-4 configurations seen in intact *ALMS1^Δex8^* cells, showing cartwheel shedding and centriole disengagement. Note the location of the cartwheel, some still in the lumen while some tethered to centrioles at the side. (**i**) UExM images of the 3-2-1 (i), 4-1-3 (ii) and 4-2-2 (iii) configurations in intact *ALMS1^Δex8^* cells showing procentriole growth. White arrows indicate newly formed procentrioles (or granddaughter centrioles). Note that CNTROB labeling marked both daughter and granddaughter procentrioles (magenta) in one of the 4-2-2 cells. All DNA was shown in blue (DAPI). All scale bars, 1μm (white) and 5μm (yellow). (**j**) Representative images showing cartwheel shedding in concert with centriole disengagement (top) and procentriole growth at abnormal locations (bottom) in *ALMS1^Δex8^* cells arrested at G2. Note the abnormal location of the cartwheel or procentriole below the proximal end of the host centriole. (**k**) Newly formed, short procentrioles in G2-arrested *ALMS1^Δex8^*cells were quantified for their locations (left) and orientations (right) relative to the host/mother centriole. Duplicated centrioles in G2-arrested WT cells were used as the control (ctrl). The relative location of each procentriole was marked on the Y axis using the length of the mother centriole as the matrix, which is normalized to one unit, with the proximal and distal ends respectively set as positions 0 and 1. Procentrioles formed below position 0 (dashed line) were frequently seen in *ALMS1^Δex8^* cells but not in WT cells. The angle between the procentriole and mother centriole was measured for each pair as the orientation. Mean ±2 s.d. from the control measurement is defined as the normal range for centriole orientation (dashed lines). Data are mean ± s.d., with samples collected evenly from three independent biological replicates. (**l**) Summary of *ALMS1^Δex8^*-induced positive feedback loops in which the cartwheel and centriole impose each other’s amplification. In brief, upon G2 arrest, the cartwheel is expanded under constraints imposed by the lumen of its own grown daughter centriole. The expanded cartwheel splits into two units, with one shedding from the lumen. The shed unit, tethered to the nearby mother centriole, grows into a procentriole, or granddaughter inside which the cartwheel is expanded again under the same constraint, while the daughter centriole that carries the other cartwheel unit disengages and drifts away, leading to the 3-x-x or 4-x-x configurations. This “expand-split-shed-grow” process occurs again at newly seeded granddaughter engaging to the mother, but not at disengaged centrioles unable to become centrosomes, leading to 5-x-x or 6-x-x configurations. Solid arrows indicate cartwheel splitting and shedding accompanying centriole disengagement, and dashed arrows indicate procentriole and cartwheel growth.

We next determined the progression of cartwheel amplification in G2-arrested *ALMS1^Δex8^* cells using a similar time-course analysis described in Fig. 1. Control (ALMS1^WT^) or mutant (*ALMS1^Δex8^*) cells synchronously entering G2 were fully blocked from mitotic entry with strong CDK1 inhibition (9.5μM RO-3306) for 0, 3, 6, 12, or 24hrs. Any non-G2 cells at the time of adding drugs were excluded by S-phase arrest (see Materials & Methods). In most control G2 cells, centrioles were found to exhibit the 2-2-0 configuration throughout the experiment (Fig. 2c). In contrast, cartwheel amplification in G2-arrested *ALMS1^Δex8^* cells was found to progress with time from 2-2-0 to 3-x-x, 4-x-x, 5-x-x, and then 6-x-x (Fig. 2d, left). By 24 hrs, >90% of cells had amplified their cartwheel, each exhibiting one of the 11 configurations (Fig. 2d, left). Intriguingly, inhibiting PLK1, a kinase whose inhibition blocks G2/M cartwheel removal (Fig. 1B), also completely blocked cartwheel amplification in G2-arrested *ALMS1^Δex8^* cells (Fig. 2d right), suggesting that these two cartwheel behaviors are related. Moreover, using live-cell imaging to trace centrioles in *ALMS1^Δex8^*cells during the time course revealed that centrioles undergo premature disengagement in G2 (Fig. 2e & Movie S1), generating disengaged centriole singlets, some of which reduplicate to form a doublet and some of which do not, shifting away from the 2-2-0 configuration. Given that all centriole singlets carry the cartwheel (Fig. S2c-p; Fig. 2b), these findings suggest that cartwheel amplification in *ALMS1^Δex8^* cells proceeds in a manner coinciding with centriole disengagement. Consistently, in cells transitioning from 2-2-0 to 3-1-2 configurations (Fig. S2d) or from 3-1-2 to 4-0-4 configurations (Fig. S2f), we observed apparent splitting of the cartwheel signal into two units.

To examine the detail, UExM was used to analyze the cartwheel during the early time points (Fig. 2f). Strikingly, unlike WT cartwheels constantly staying inside the lumen of daughter centrioles under G2 arrest, we found that *ALMS1^Δex8^* cartwheels can split and shed in concert with centriole disengagement (Fig. 2f). In cases where the daughter centriole, marked by CNTROB, was in an early phase of disengagement, as judged by its relatively close, juxtaposed positioning to the mother (Fig. 2f; #1), the cartwheel of the daughter had split into two units, with the distal unit remaining in the lumen while the proximal unit was either partially (Fig. 2f; #1) or completely (Fig. 2f; #2) shed from the lumen and tethered to the nearby mother centriole. Ex-dSTORM analyses confirmed physical splitting and shedding of the cartwheel in the centriole lumen (Fig. 2g). Notably, the shed cartwheel was frequently observed attached to the adjacent disengaging mother centriole at irregular positions or angles (Fig. 2f), a pattern contrasting with the typical growth of new cartwheels, which consistently occurs at the sides of mother centrioles in orthogonal arrangements (refer to Figs. 2j & k below for further details).

In cases where centriole disengagement had led to a farther separation of the daughter from the mother (Fig. 2f; #3), we found that the shed cartwheel remained tethered to the mother, whereas the disengaged daughter kept the other cartwheel unit in the lumen (Fig. 2f; #3). The combination of cartwheel shedding and centriole disengagement transforms the 2-2-0 configuration in a typical G2 cell to either the 3-1-2 or 4-0-4 configuration, which is precisely what we observed among cells that carried 3 cartwheels and 4 centrioles or 4 cartwheels and 4 centrioles, respectively, without exception (Fig. 2h). Moreover, the growth of the shed cartwheel, which is tethered to the mother, into a procentriole further transforms the 3-1-2 to 3-2-1 configuration (3 cartwheels and 5 centrioles) or the 4-0-4 to 4-1-3 (4 cartwheels and 5 centrioles) and 4-2-2 (4 cartwheels and 6 centrioles) (Fig. 2i; arrows indicate procentrioles). Newly formed procentrioles undergoing the same cartwheel amplification would lead to configurations with 5 or 6 cartwheels (5-x-x or 6-x-x), which is consistent with what we observed (Fig. S2c-p). For example, cartwheel shedding from a granddaughter or great-granddaughter can cause a shift from the 4-1-3 to 5-0-5 (Fig. S2q; see split cartwheel #1 and #2) or 5-2-3 to 6-1-5 configuration (Fig. S2s; see split cartwheel #2 and #3), respectively. For the 5-2-3 (Fig. S2r) or 6-2-4 configuration (Fig. S2t), two cartwheels growing into procentrioles were seen at the side of mother centrioles, whereas the other four cartwheels remained in the lumen of disengaged centriole singlets. Furthermore, by delineating the nascent, short procentriole tethered to the host centriole regarding its position and orientation (Fig. 2j & k), we observed that a considerable proportion of these procentrioles are either developing at atypical sites, such as beneath the proximal end of the host centriole (Fig. 2k, left), or are oriented at angles beyond the typical range established by wild-type centrioles (Fig. 2k, right), which aligns with the stochastic nature of cartwheel shedding-tethering during centriole disengagement. These highly patterned phenotypes suggest that ALMS1 mutations can induce cartwheel splitting and shedding during G2 arrest, coinciding with centriole disengagement, resulting in positive feedback loops where centrioles and cartwheels amplify each other (see Fig. 2l legend; Discussion for summary).

### ALMS1 is in complexes with CEP152, CEP63, or PCNT, both independently and outside of the cartwheel and centriole

ALMS1 is involved in S-phase cartwheel formation (Fig. 1i; Fig. S1h & j) and G2/M cartwheel dynamics (Fig. 2l), suggesting that it may interact with cartwheel assembly factors. Immunoprecipitation (IP) of the endogenous ALMS1 from clarified cell extracts did not co-immunoprecipitate cartwheel components like PLK4 and SAS-6 (not shown) but did pull down pericentrin (PCNT), CEP152, and CEP63, which is known to interact with CEP152^16–18^ (Fig. 3a, left). It has been reported that CEP63 is hardly detectable by Western blot unless it is immunoprecipitated^17^. Indeed, immunoprecipitating the endogenous CEP63 pulled down not only CEP152 but also ALMS1 and PCNT (Fig. 3a, left). Interestingly, however, UExM analyses showed that ALMS1 is not co-localized with CEP152, CEP63, or PCNT at centrioles (Fig. 3b), implying that the association of ALMS1 with these molecules occurs outside the centriole. Similar Co-IP results were seen with cell extracts prepared from *STIL^-/-^* cells lacking the cartwheel (Fig. 3a, right), further suggesting that ALMS1 can form soluble complexes with CEP152, CEP63, or PCNT independently of the cartwheel and centriole. Given that PCNT is known to mark an amorphous cloud in the cytoplasm from which *de novo* centrioles form in multiciliated cells^33^, and that CEP152 and CEP63 form well-characterized complexes that recruit PLK4 for activating cartwheel assembly^15–18^, our findings support the idea that ALMS1 promotes centriole biogenesis at a step prior to cartwheel assembly.

**Fig. 3.**
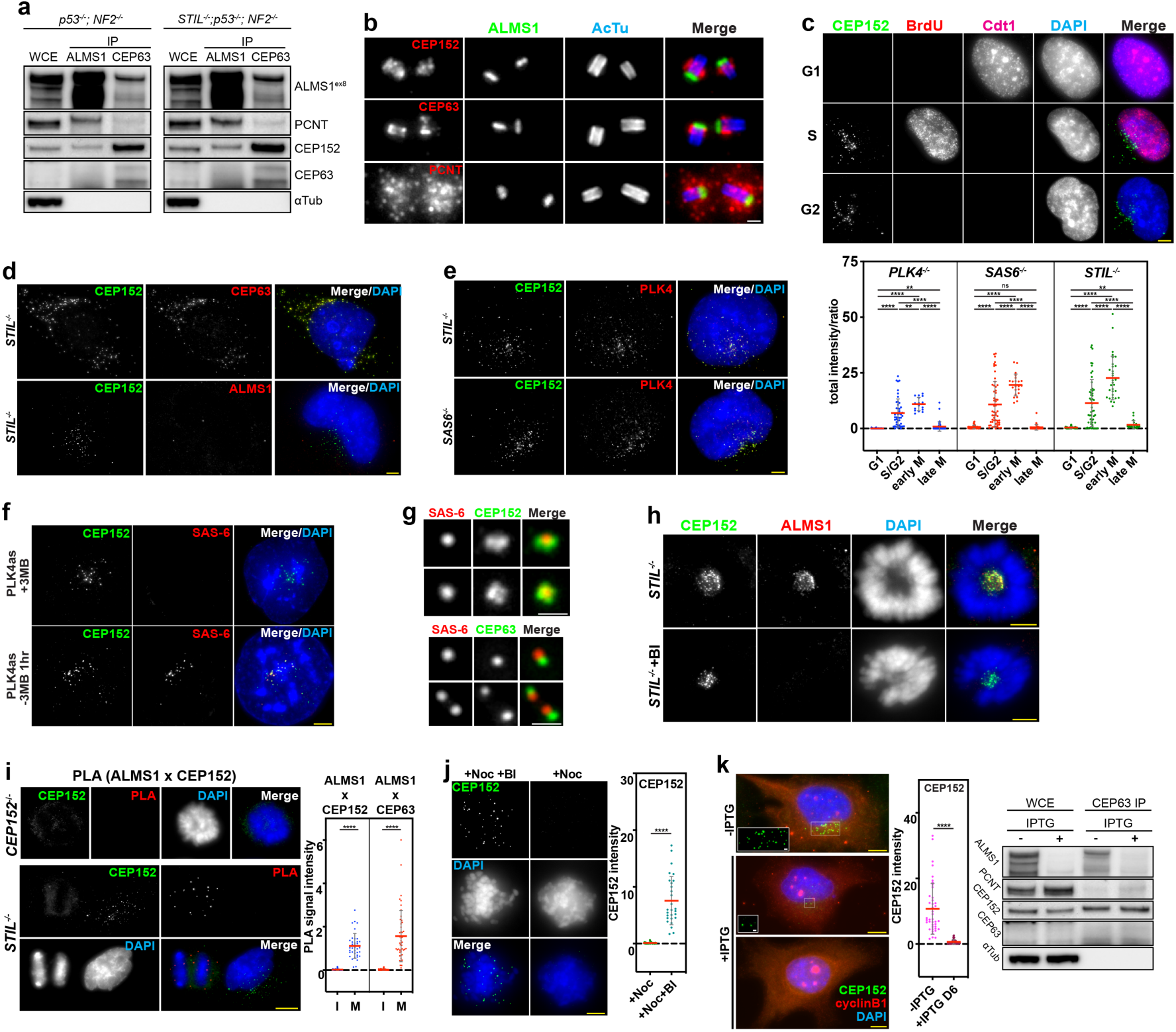
ALMS1 mediates the cyclical assembly-disassembly dynamics of CEP152-CEP63-PCNT composites that function as the seed for cartwheel formation. (**a**) Immunoprecipitation (IP) of the endogenous ALMS1 or CEP63 from either control cells (*NF2^-/-^ ; p53^-/-^*) or cartwheel-KO cells (*STIL^-/-^*). Antibodies used for IP and western blot were indicated. Note that before IP, the endogenous CEP63 in the whole cell extract (WCE) was undetectable, as reported previously. (**b**) UExM images of mature centrioles showing non-colocalizations of ALMS1 with CEP152, CEP63, or PCNT. (**c**) CEP152 puncta formed in S or G2 and unformed during mitosis. Quantification of CEP152 puncta in indicated cartwheel-KO cell lines. Cdt1 indicates G1, BrdU marks S, and G2 cells lack both. (**d**) CEP152 formed composite structures with CEP63 but not with ALMS1. Images shown here were from *STIL^-/-^* cells. Other co-localization studies were shown in the Supplemental Figure S3b. (**e**) CEP152 composites in *STIL^-/-^* or *SAS-6^-/-^* cells can robustly recruit PLK4. (**f**) CEP152 composites drive cartwheel or SAS-6 assembly. *PLK4^as^* cells inducibly expressing analog-sensitive PLK4 in the presence of 3MBPP1 form CEP152 composites (top panel). Upon washout of 3MBPP1 for 1 hr, the cartwheel signal emerges from the CEP152 composite (bottom panel). (**g**) UExM images of SAS-6 assembly initiated from the CEP152 (top) or CEP63 puncta (bottom) 1 hr after 3MBPP1 washout as described in (**f**). (**h**) ALMS1 is timely recruited to CEP152 composites upon mitotic entry, prior to their disassembly, in a PLK1-dependent manner. Asynchronously growing *STIL^-/-^* cells treated with or without PLK1 inhibition (200 nM BI-2536) for 6 hours were fixed. Cells just entering mitosis in their early prophase were examined for CEP152 and ALMS1 localization. Additional images revealing timely recruitment of ALMS1 during G2/M were shown in Supplemental Figure S3c. (**i**) PLA reactions detecting the interaction of ALMS1 with CEP152 or CEP63 in interphase and mitotic cells. A representative image of PLA reactions that detect interactions between ALMS1 and CEP152 (ALMS1 x CEP152) in a pair of mitotic and interphase *STIL^-/-^* cells that co-stained with CEP152 composites. *CEP152^-/-^*cells were used as the negative control for PLA reactions. PLA signals for ALMS1 interaction with CEP152 or CEP63 in interphase and mitosis were quantified (right). See also the Supplemental Figure S3d. (**j**) The disassembly of CEP152 composites or CSs in mitosis requires PLK1 activities. Interphase *SAS-6^-/-^* cells obtained by shaking off mitotic cells were allowed to enter and arrest in mitosis in the presence of nocodazole alone (+Noc) or nocodazole and BI-2536 (+Noc +BI) for 6 hours. Mitosis-arrested cells were then examined. (**k**) Depletion of ALMS1 abolished CS assembly without impacting the level of CS components CEP152, CEP63, and PCNT. *SAS-6^-/-^*cells stably introduced with IPTG-inducible shRNA targeting ALMS1 were generated and used. Six days after IPTG induction, CS assembly was examined and quantified in S or G2 cells, as marked by cyclin B1 expression. CS components before and after ALMS1 knockdown were analyzed by western blot (right). CEP63 was detected after IP. All DNA was shown in blue (DAPI). All scale bars are 1 μm (white) and 5 μm (yellow). Statistical analyses (**c,** & **i-k**) were done by Welch’s t-test (p < 0.01).

### Composite objects comprising CEP152, CEP63, and PCNT but not ALMS1 form independently of the cartwheel during S or G2 phase, which are disassembled in mitosis like the cartwheel

To explore this idea, we examined the cell cycle dynamics of CEP152 in *PLK4^-/-^, STIL^-/-^*, or *SAS-6^-/-^*cells, where cartwheel assembly is knocked out. In all cases, we found that CEP152 forms cytoplasmic puncta in highly variable numbers during S and G2 phases (Fig. 3c; Fig. S3a), which robustly disappear during mitosis (Fig. 3c; Fig. S3a), a pattern resembling that of the cartwheel dynamics. Immunostaining further showed that CEP152 puncta are composite structures containing CEP63 and PCNT (Fig. 3d; Fig. S3b) but, surprisingly, not ALMS1 (Fig. 3d; Fig. S3b), which we hereafter called CEP152, CEP152-CEP63, or CEP152-CEP63-PCNT composites. This finding suggests that ALMS1 does not form macromolecular composite structures with its interacting partners, neither the CEP152 composite formed in the absence of centrioles nor those recruited to mature centrioles (Fig. 3b).

### CEP152-CEP63-PCNT composites function as the seed for cartwheel formation, hereafter called the "cartwheel seed" or "CS"

To explore the function of CEP152-CEP63-PCNT composites, we first examined if they recruit PLK4 in *STIL^-/-^* or *SAS-6^-/-^* cells where cartwheel assembly is knocked out but PLK4 is present. We found that PLK4 is robustly recruited to nearly all composites (Fig. 3e). To see if these PLK4-containing composites can drive cartwheel formation, we used *PLK4^as^*(analog-sensitive) cells in which the endogenous PLK4 is deleted and replaced with a mutant PLK4 sensitive to bulky ATP analogs 3MBPP1 and whose expression is under the doxycycline (DOX) inducible promoter^14^. Like *STIL^-/-^*or *SAS-6^-/-^* cells, *PLK4^as^* cells arrested at S phase in the presence of 3MBPP1 and DOX were found to form variable numbers of CEP152 composites (Fig. 3f). Intriguingly, 1hr after washing out 3MBPP1, we observed robust cartwheel signals (SAS-6) emerging from most of the CEP152 composites (Fig. 3f). UExM revealed that the cartwheel appears to locate at a signal-sparse area, or cavity, of the CEP152 structure (Fig. 3g) or juxtaposed to the CEP63 structure (Fig. 3g). These results suggest that CEP152 and CEP63 form structured composites capable of driving cartwheel assembly independently of centrioles, acting like the seed. Moreover, the presence of variable numbers of CEP152-CEP63-PCNT composites per cell explains how centrioles that form in the absence of preexisting centrioles are highly variable in numbers. For simplicity, we hereafter used the term “cartwheel seed” or “CS” interchangeably with “CEP152 composite” or “CEP152-CEP63-PCNT composite” to describe this novel function.

### ALMS1 mediates the cyclical assembly-disassembly dynamics of the CS

ALMS1 is linked to G2/M cartwheel dynamics in localization (Fig. 1h) and function (Fig. 2l). Given that CEP152-CEP63-PCNT composites, which function as CSs, also disappear during mitosis (Fig. 3c) like the cartwheel, we asked if ALMS1 is physically connected to CS disassembly. Immunofluorescence microscopy revealed that while ALMS1 is not part of CSs during S/G2 (Fig. 3d; Fig. S3b & c), it is recruited to CSs at late G2 or early M, prior to CS disassembly (Fig. 3h; Fig. S3c), in a PLK1-dependent manner (Fig. 3h; Fig. S3c). To test if ALMS1 interacts with the components dissociating from the CS, the proximity ligation assay (PLA) was used to visualize single protein-protein interactions in fixed cells^34, 35^. Indeed, detecting complex formation of ALMS1 with CEP63 or CEP152 revealed robust PLA signals in mitosis when CSs disappear but not in S or G2 phase when CSs persist (Fig. 3i; Fig. S3d). Moreover, like cartwheel removal (Fig. 1b), we found that CS disassembly in mitosis requires PLK1 (Fig. 3j), further supporting that cartwheel dynamics are regulated through CS dynamics. To explore the function of ALMS1-mediated CS dynamics, we generated *SAS-6^-/-^* cells stably expressing doxycycline-inducible shRNA constructs targeting ALMS1. Intriguingly, 6 days after inducing ALMS1 knockdown in *SAS-6^-/-^* cells, CS formation in S or G2 cells, as marked by cyclin b1 expression, is nearly all abolished (Fig. 3k), consistent with ALMS1 being required for cartwheel and centriole formation altogether (Fig. 1i; Figs. S1h & j). Furthermore, we found that ALMS1 depletion does not impact the cellular level of the CEP152-CEP63 complex or PCNT (Fig. 3k; right), suggesting that ALMS1 is involved in transforming CEP152, CEP63, and PCNT into CSs, rather than stabilizing them. Our findings together show that vertebrate centriole biogenesis is grounded on ALMS1-mediated, cyclical CS assembly-disassembly dynamics, with ALMS1 acting as an external cofactor not incorporated into the CS, cartwheel, or procentriole.

### Ex-dSTORM delineates a nanoscale ring-like organization of CEP152 and CEP63 in the CS

We next explored the physical or structural properties of CSs. Sucrose gradient fractionation analyses revealed that CSs can be purified under a similar condition used for centrosome isolation, which are enriched at the interface between 50% and 70% sucrose solutions (Fig. 4a), revealing CSs as dense objects. UExM analyses of CSs in fixed cells showed that CEP152 forms a ring-like structure of which CEP63 is localized at the center (Fig. 4b; Movie S2). PLK4 molecules recruited to CSs were also found to surround CEP63 in ring-like patterns (Fig. 4B). Intriguingly, by analyzing the CS with Ex-dSTORM that achieves true molecular resolution (sub-5 nm), we found that CEP63 is resolved into a ring of approximately 58 nm in mean diameter (Fig. 4c-e), which is encircled by a larger CEP152 ring (Fig. 4d) averaging about 136 nm in diameter (Fig. 4e). We often observed incomplete rings with gaps, missing one or more CEP152 or CEP63 dots (Fig. 4e & 4f), which may reflect intrinsic CS dynamics, or alternatively, incomplete labeling of the CS under Ex-dSTORM due to technical reasons. Analyzing the ring pattern revealed that the average angular intervals between CEP152 or CEP63 dots in each ring appear to be multiples of ∼40 degrees, inferring 8-, 9-, or 10-fold radial symmetry (Fig. 4f & g). Moreover, from the side view, CEP63 rings could be seen as single units, as a stack of two units, or as two closely interacting structures (Fig. 4h), implying dynamic arrangements of CSs. Note that under cartwheel-knockout conditions, CSs are unable to develop into static structures like centrioles. Together, CSs are dense, nanoscale, concentric ring-like structures capable of seeding cartwheel formation.

**Fig. 4.**
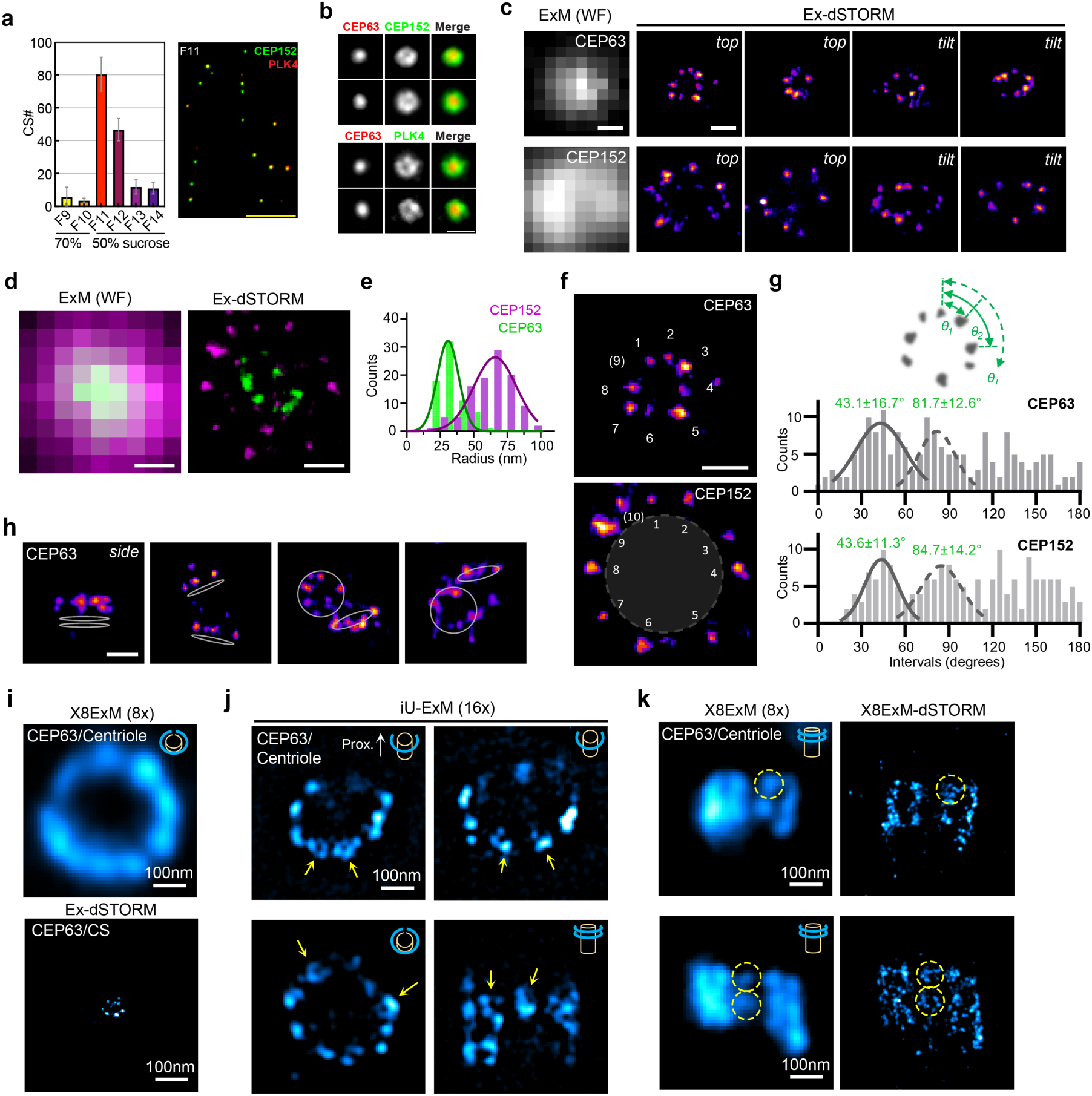
The CS is a dense, structured composite with CEP152 and CEP63 forming a nanoscale ring-like organization. (**a**) Sucrose gradient fractionation of CSs from S-phase arrested *STIL^-/-^* cells. CSs were peaked at the fraction (F11) collected from the interface between 70% and 50% sucrose solutions (left). Purified CSs from F11 were pelleted onto coverslips and examined with indicated antibodies (right). Scale bars, 10 μm. (**b**) UExM images of CSs in fixed *STIL^-/-^* or *SAS-6^-/-^* cells labeled with CEP152, CEP63, or PLK4 as indicated. Scale bars, 1 μm. (**c**) Representative dSTORM images of expanded CSs in cells labeled with CEP63 or CEP152, in top and tilt views. Ex-dSTORM resolves the spatial distributions of CEP63 and CEP152 at sub-5 nm resolution. Scale bars, 50 nm. (**d**) Dual-color Ex-dSTORM images of CSs co-labeled with CEP63 and CEP152, showing a concentric spatial arrangement. Scale bars, 50 nm. (**e**) Quantifications revealed that CEP63 and CEP152 form rings with radii of 29 ± 8 nm and 68 ± 16 nm, respectively. Data are presented as mean ± s.d., based on 69 and 106 measurements from more than 9 CSs each. (**f, g**) Angular spacing analysis of CEP152 and CEP63 dots within individual rings, with angular intervals averaging in multiples of ∼40°. Data are presented as mean ± s.d.. Scale bars, 50 nm. (**h**) Side-view Ex-dSTORM images revealing CSs (CEP63 rings) existing as a stack of two units (1^st^ panel, left), single individual units (2^nd^ panel), or two close-by structures interacting with each other (3^rd^ & 4^th^ panels), implying dynamic CS configurations under the cartwheel-KO condition. Scale bars, 50 nm. (**i**) UExM images at 8-fold expansion (X8ExM) of CEP63 either at the centriole (top) or at the CS (bottom) under the same scale bar (100 nm). Cartoons show the cross-section of the centriole (yellow) and CEP63 (green). (**j**) iU-ExM images at 16-fold expansion revealed some CEP63 puncta at centrioles as a ring (arrows). Cartoons show the orientation of the centriole (yellow cylinders) and associated CEP63 (green rings), with proximal ends pointing upward. Centrioles encircled by one or two layers of CEP63 were shown. (**k**) X8ExM-dSTORM images at ∼2 nm resolution that resolve CEP63 puncta into a ring-like structure resembling the CS (circles). Centrioles encircled by two or more layers of CEP63 puncta were shown.

### CSs at canonical centrioles

Centriole-associated CEP152/CEP63 complexes are known to recruit PLK4 for canonical centriole duplication^15–18^. By modeling the centriole, CEP152 or CEP63 puncta were reported to collectively organize into a large, centriole-encircling ring of ∼300 nm in diameter^15, 36^, a pattern we confirmed here and found to be categorically distinct from the CS in both size and context (Fig. 4i). Intriguingly, further analyses revealed that some of these CEP63 puncta at mature centrioles can be individually resolved into ring-like structures resembling CSs in size, either by iterative ultrastructure expansion microscopy (iU-EXM) that achieves ∼16x molecular expansion^37^ (Fig. 4j) or by Ex-dSTORM (Fig. 4k), supporting the idea that canonical centriole duplication also involves CSs or CS-like structures. We hypothesize that each CEP63 punctum at mature centrioles is potentially a CS or precursor unit and that for number control during canonical duplication, only one unit is mature or active for cartwheel formation.

### ALMS1 depletion and addback initializes centriole biogenesis anew with diverse yet heritable architectures that evolve via selection, instead of restoring canonical assembly

ALMS1 drives CS formation independently of the centriole-cartwheel relationship, suggesting that it contains or mediates the initial transformation cues for centriole biogenesis outside the cartwheel and centriole. If the cue is innate to ALMS1, centriole biogenesis should be recoverable and remain unchanged after ALMS1 depletion and add-back, like what has been reported previously with PLK4 depletion and add-back^38^. As the control, we found that inactivation and restoration of PLK4, which interrupts the cartwheel but not CS assembly cycle (Fig. 3c), indeed leads to the generation of supernumerary centrioles with the same canonical shape and size (Fig. 5a). In contrast, using inducible shRNA to deplete and restore ALMS1, which interrupts CS assembly cycles, produced *de novo* centrioles with striking diversity, including irregularities in shape and size, with multiple centrioles of distinct architectures coexisting within the same cell (Figs. 5b, S4a). Beyond morphology, we observed thin centrioles lacking γ-tubulin accumulation in the lumen (Fig. S4b), unusually short centrioles duplicating without cartwheel removal (Fig. S4c), and distinct forms of centrioles with dramatically different PCM sizes undergoing duplication in the same cell (Fig. 5c), all previously unreported. The formation of centrioles with heterogeneous architectures within the same cell—rather than with either uniformly normal or uniformly defective features—challenges the view that regards centriole assembly as deterministic by genetics. Instead, it argues that perhaps the initial cue preserved by ALMS1 is adaptable.

**Fig. 5.**
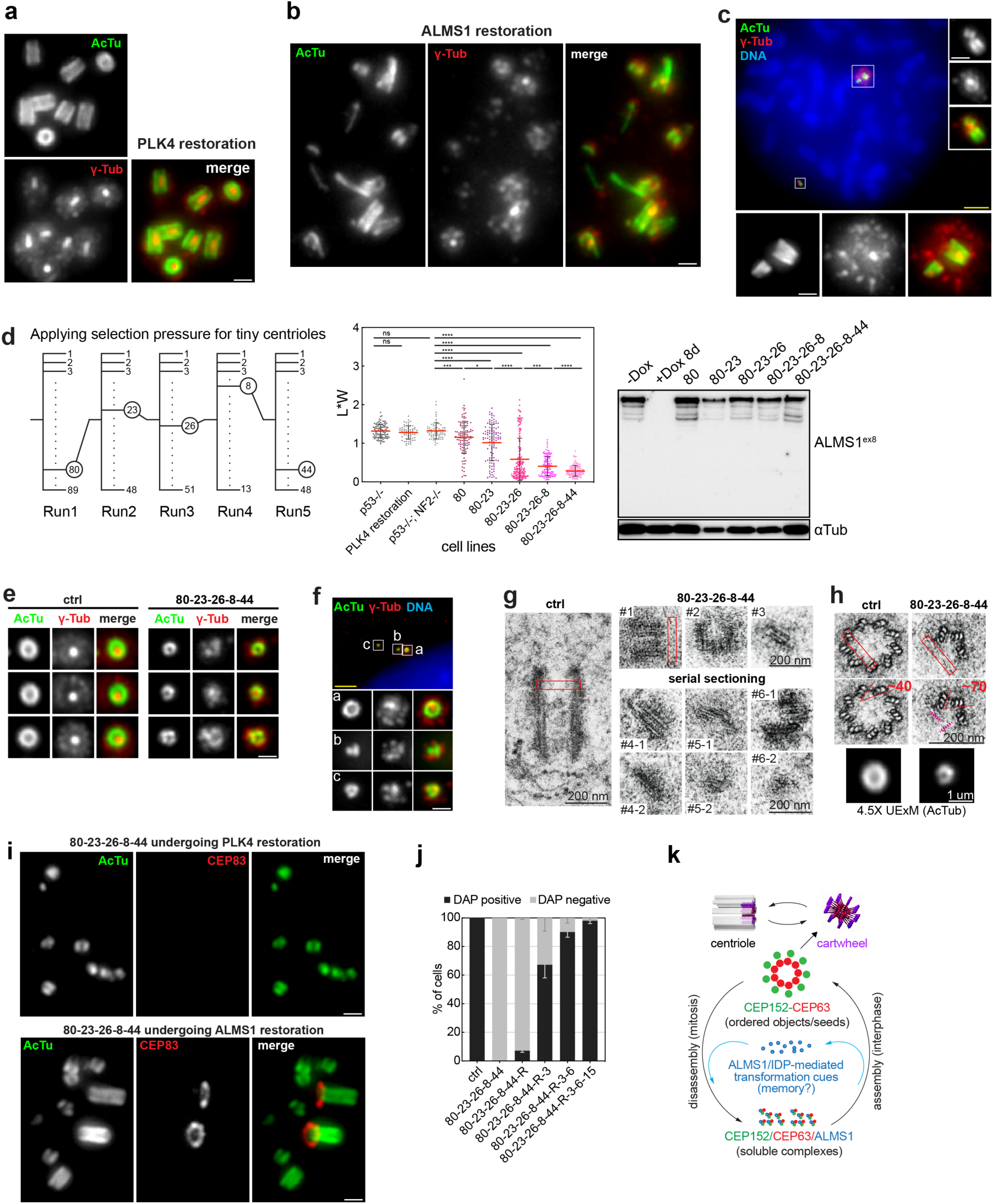
ALMS1 depletion and addback initializes centriole biogenesis anew, creating diverse yet heritable architectures that evolve through selection. (**a**) Canonical centrioles formed in variable numbers upon PLK4 inhibition and reactivation (PLK4 restoration). *NF2^-/-^; p53^-/-^*cells were treated with the PLK4 inhibitor Centrinone for 10 days, followed by drug washout for 5 days. (**b**) Centrioles with diverse shapes or sizes emerged in the same cell upon ALMS1 depletion and addback (ALMS1 restoration) by inducible shRNA. Stable clones of *NF2^-/-^; p53^-/-^* cells carrying two DOX-inducible shRNA constructs targeting ALMS1 were treated with DOX for 7 days and reseeded in the absence of DOX for 5 days. (**c**) Two pairs of duplicated centrioles with extreme differences in size co-existing in early prophase. Note the size difference in centrioles reflects on the size of the surrounding PCM (γ-tubulin; red). (**d**) Applying selection pressure for tiny centrioles through a series of clonal cell isolations. Left, a diagram summarizing 5 runs of clonal cell isolation, each including examination time, taking 6-8 weeks. In each run, the clone with the highest frequency of forming tiny centrioles was selected (circle). Middle, quantification of the centriole size in each selected clone, inferred by the length and width (L*W). After 5 runs, cells in clone 80-23-26-8-44 grow a heterogeneous group of tiny centrioles whose sizes do not overlap with that of WT centrioles (Welch’s t-test, p < 0.01). Right, ALMS1 levels were restored in each isolated clone, determined by Western blot, with α-tubulin as the loading control. (**e**) Representative UExM images of tiny centrioles from clone 80-23-26-8-44 cells that form partial cylinder shapes, the diameter of which is distinct from that of control centrioles. (**f**) UExM images of 3 tiny centrioles with different shapes co-existing in the same cell from clone 80-23-26-8-44. (**g** and **h**) Transmission electron microscopy (TEM) analyses of control or tiny centrioles. (**g**), One control and six tiny centrioles (#1-#6) were shown, including serial sectioning of three tiny centrioles (#4-#6). #4 tiny centriole appears to have 3 microtubule blades in two consecutive sections. Red boxes marked the width of WT centrioles for comparison. (**h**), One control and one tiny centriole in cross section were shown. Note that the tiny centriole has three microtubule triplets arranged at a ∼70-degree angle instead of the ∼40-degree angle seen for control centrioles. Red boxes marked the width of WT centrioles. UExM images of WT and tiny centrioles from (**e**) were placed below the EM image for easy comparison. (**i**) Clone 80-23-26-8-44 cells going through PLK4 (top) or ALMS1 (bottom) restoration were examined for *de novo* centrioles. Larger centrioles capable of recruiting appendages formed and co-existed with tiny centrioles 5-6 days after ALMS1 restoration but not PLK4 restoration. (**j**) Quantification of 80-23-26-8-44 cells hosting appendage-equipped centrioles before and after ALMS1 restoration (80-23-26-8-44-R). After 3 runs of clonal cell isolation following ALMS1 restoration, a clone (80-23-26-8-44-R-3-6-15) where >95% of cells carry appendage-equipped centrioles was isolated. All DNA was shown in blue (DAPI). All scale bars, 1 μm (white) and 5 μm (yellow). (**k**) A model summarizing (i) the cartwheel-centriole assembly cycle seen under the *ALMS1^Δex8^* condition and (ii) the ALMS1-mediated CS assembly-disassembly cycle. Each of the two can drive reproducible centriole biogenesis, with ALMS1 acting extrinsically without incorporating into the CS, cartwheel, or procentriole. As ALMS1 depletion and addback cause de novo centriole biogenesis with diverse yet heritable architectures, we hypothesize that ALMS1, an intrinsically disordered protein, may store and transmit molecular memory as the transformation cue for adaptable centriole biogenesis (see the Discussion).

To determine if centrioles with diverse architectures that form following ALMS1 depletion-addback are transient, nonspecific byproducts or, alternatively, stable structures capable of propagation, we traced them via a series of clonal cell isolations (Fig. 5d). 7 days after ALMS1 restoration, single cells each carrying supernumerary centrioles varying in shape or size were clonally seeded in 96-well plates and grown for 4 weeks, during which centrioles propagated and segregated randomly into descendant cells. Cells from each clone were examined by UExM for the frequencies at which centrioles of various sizes were maintained. Our selection process focused on extremely small centrioles, here called tiny centrioles, because they stain weakly with markers like centrin or γ-tubulin, making them distinguishable from normal centrioles even under regular epifluorescence microscopy (not shown). Cells in clones that carried tiny centrioles at higher frequencies were selected for further propagation through additional runs of subclonal cell isolations, a process mimicking natural selection. Note that these tiny centrioles had coexisted with larger centrioles in the same cell from the beginning and that their segregation during cell division is a stochastic process on which our selection pressure applies. Intriguingly, after 5 runs of clonal and subclonal cell isolation, a lineage of descending cell lines was obtained (Fig. 5d), each carrying progressively higher frequencies of tiny centrioles while expressing similar levels of ALMS1 (Fig. 5d). Clones carrying higher frequencies of abnormally large centrioles were also seen but not followed here. Clone 80-26-23-8-44 was found to stably grow a heterogenous group of thin or short centrioles not overlapping with the size of wild-type centrioles (Fig. 5d). Immunostaining revealed that these tiny centrioles can recruit most centrosomal markers we examined (Fig. S4d), except for distal and sub-distal appendages, which were completely absent (0%, *n* = 300) (Fig. S4e). Moreover, most of these tiny centrioles either failed to form a cylinder shape (Figs. S4d & S4e) or formed a partial cylinder with variable diameters (Fig. 5e), which were coexisting in the same cells (Fig. 5f). Transmission electron microscopy (TEM) confirmed the UExM result (Figs. 5g & h), showing small centrioles with fewer sets of microtubule blades (Fig. 5g) or partial cylinders with microtubule triplets arranged at ∼70-degree angles from each other (Fig. 5h), instead of ∼40-degree angles for control centrioles.

To check if tiny centrioles can form reproducibly in the absence of preexisting copies, clone 80-26-23-8-44 cells were allowed to undergo PLK4 inhibition and reactivation. Indeed, all centrioles formed after PLK4 reactivation were found to be small and without appendages (0%, *n* = 300) (Fig. 5i; top). To explore the adaptability or plasticity of centriole biogenesis in the presence of reproducibility, we asked if clone 80-26-23-8-44 cells with tiny centrioles can be re-initialized to form larger centrioles via another round of ALMS1 depletion and addback. Intriguingly, after ALMS1 restoration, larger centrioles capable of recruiting appendages were seen randomly emerging together with tiny centrioles in the same cell (Fig. 5i; bottom). Moreover, under the selection pressure, these newly formed appendage-bearing centrioles could outcompete tiny centrioles after 3 rounds of clonal cell isolations, propagating reproducibly (Fig. 5j, 80-26-23-8-44-R-3-6-15). In short, we found that—while ALMS1 is extrinsic to the CS or cartwheel—centriole biogenesis fails to initiate without it, whereas reintroducing ALMS1 creates diverse yet heritable characteristics that evolve through selection. Together, our findings prompt a conjecture that centriole biogenesis may fundamentally involve memory (Fig. 5k; see the Discussion below).

## Discussion

How orderless constituents are transformed into cartwheels or centrioles has not been known in any condition. Our results here show that, under *ALMS1^Δex8^* mutant conditions, the centriole and cartwheel can drive or impose each other’s amplification through a feedback loop (Fig. 2l & 5k): (i) an existing cartwheel forms a centriole and becomes embedded in the centriole lumen. (ii) The embedded cartwheel is expanded under constraints imposed by the lumen of its own grown centriole. (iii) The expanded cartwheel splits and sheds. (iv) The shed cartwheel is then free to grow into a new centriole with the identical shape, starting the next cycle. This complementary, cartwheel-centriole relationship can be maintained via continuous cycles wherein outputs become the next inputs—like memory transfer—without invoking a single initiating cause. Centriole biogenesis that naturally occurs via this complementary relationship has not been reported or carefully looked for in organisms with compatible lifestyles, but it is clearly incompatible with organisms like vertebrates, where centriole elimination and regeneration occur under physiological conditions. To accommodate such centriole turnover, are there transformation cues propagating beyond the centriole-cartwheel relationship?

Our studies here identify a dense, ring-like CEP152-CEP63 composite of ∼60 nm in diameter that acts as the cartwheel seed (CS) propagating independently of the cartwheel-centriole relationship via ALMS1, preserving the initial transformation cue for biogenesis (Fig. 5k). We show that ALMS1 is an external cofactor functionally linking CS assembly and disassembly into a cyclical process without itself incorporating into the CS. Erasing the old cycle followed by reforming a new cycle through ALMS1 depletion and addback allows centrioles with diverse yet heritable shapes to form and propagate, revealing adaptability and reproducibility—often viewed as mutually exclusive—as the co-feature. To contemplate how the “dual feature” may work, we hypothesize, in general terms, that ALMS1 may possess the capacity to store (write) and retrieve (read) molecular memories. ALMS1 is a large intrinsically disordered protein (IDP), with >95% of its 4169 amino acid residues predicted to lack stable secondary structure^24^. IDPs are known to adopt and convey specialized shapes through context-dependent protein-protein interactions^39, 40^, a property consistent with ALMS1’s role in mediating adaptability and reproducibility of centriole formation (Fig. 5). We imagine, for instance, that by interacting with existing CSs during CS disassembly (Fig. 3), naïve ALMS1 adopts information (memory), and by interacting with naïve CEP152/CEP63 complexes, the adopted ALMS1 conveys that memory for CS assembly—a loop where information is adjustable yet transmittable, allowing heritable phenotypic variations to form and adapt without entailing random, often detrimental, genetic mutations.

## Methods

### Cell culture

Human telomerase-immortalized retinal pigment epithelial (RPE1) cells were cultured in DME/F-12 (1:1) medium supplemented with 10% FBS and 1% penicillin-streptomycin. U2OS cells were cultured in McCoy’s 5A medium with 10% FBS and 1% penicillin-streptomycin. BrdU was added to culture medium at 30 μM to label S-phase cells. Aphidicolin (A0781, Sigma) was used at 2 μg/ml to arrest cells in S phase. CDK1 inhibitor, RO-3306 ^41^ (217699; Sigma), was used at 9.5 μM to fully arrest cells at a point before the G2/M boundary. BI-2536 was used at 200 nM to inhibit PLK1. MG132 was used at 10 μM to inhibit the proteasome. Centrinone (HY-18682, MCE) was used at 125 nM to inhibit PLK4. Cells in which the endogenous PLK4 was replaced with the PLK4 analog-sensitive mutant were treated with 2 μM 3MB-PP1 (529582, Sigma) to eliminate centrosomes.

### RNAi

Transient transfection of siRNA oligos was performed using lipofectamine RNAiMAX (Thermo Fisher). The sequences of ALMS1 siRNA oligos are GCCGAUUGUUAACUACAAA and GCACGAAAGGAGUAGCUCU (s15391 & s15392; Thermo Fisher), both working effectively. The control siRNA oligo is the Silencer® Select Negative Control No. 1 (Thermo Fisher). The targeting sequences of lentiviral shRNA constructs that constitutively knock down ALMS1 expression include TTGTGAGTCTGGTTGAATAAA (3’-UTR), CGGGTGTATCTAATGGTGATT (exon 8), and TATTGGCACACAGACGAATTT (exon 8), all commercially available from Sigma-Aldridge (TRCN0000424420, TRCN0000083999, and TRCN0000428950, respectively). For inducible ALMS1 knockdown, two equally effective systems were established: (i) the IPTG inducible shRNA construct inserted into the AAVS1 locus by CRISPR knock-in and expressing the highly effective 3’-UTR targeting sequence TTGTGAGTCTGGTTGAATAAA, and (ii) a combination of two doxycycline inducible lentiviral constructs, TGAAATATCACTAATATCTCTG (exon 8) and TATATAATTTAATTCGTCTGGG (exon 16), both designed with the SplashRNA algorithm^42^ and expressed from miR-E backbone vectors^43^. Cell lines stably expressing doxycycline or IPTG inducible hairpins for ALMS1 knockdown were clonally isolated and maintained.

### Genome editing

Genetic inactivation of the ALMS1, NF2, p53, CEP152, SAS-6, or STIL gene was created by CRISPR/Cas9-mediated gene targeting^44^. The *CEP295^-/^*^-^ cell line was previously published^20^. The PITCh genome integration method^45^ was used to endogenously tag ALMS1 molecules with human influenza hemagglutinin (HA) epitopes at the N-terminus. The following gRNAs were used in this study, GGCGAACGTGGACGACGTAG (for ALMS1_exon1 knockout), GGACTAGCTGACCAGACAAC (for ALMS1_exon8 knockout), CCCAGAGCGAGACACCAACA (for ALMS1 tagging), CCTGGCTTCTTACGCCGTCC (for NF2 knockout), and GGGCAGCTACGGTTTCCGTCTGG (for p53 knockout). TGTCATTAGACTTTGGCAGT (for CEP152 knockout), GTGAAATGCAAAGACTGTG (for SAS-6 knockout), and TGTGGAATTTGACTTGCAT (for STIL knockout).

### Preparation of a bulk of cells synchronously entering G2

The CDK4/6 inhibitor palbociclib has been used to synchronize the growth of RPE1 cells from G1 to M^26^. In this published protocol, the cell-cell contact inhibition known to impact the synchrony of cell cycle progression is minimized by growing cells at very low densities, a condition, however, not ideal for our UExM analyses of centrioles^26^. We thus genetically disrupted contact inhibition in RPE1 cells by inactivating the NF2 gene, generating *NF2^-/-^; p53^-/-^ or NF2^-/-^; p53^-/-^; ALMS1^Δex8^* cell lines. Cells lacking NF2 were seeded at 25,000 cells/cm² and grown in the presence of 200nM of Palbociclib (S1579, Selleckchem) for 36 hours, when most of the cells were arrested in G1 at the restriction point^26^. By washing out palbociclib, cells were released from the restriction point and started to progress synchronously to mitosis, taking about 12-14 hours^26^. Around 12 hrs after the release, when a bulk of cells were in G2, different amounts of the CDK1 inhibitor RO-3306 were added to the culture, with a goal to either fully block mitotic entry (9.5μM) or to slow down the G2/M transition (6μM). For time-course experiments, which require pure G2 populations of cells, aphidicolin (2μg/mL) was added together with RO-3306 to block any cells not yet reached G2 at the time of the experiment from ever entering G2. Cells arrested at G2/M can be easily distinguished from S-phase cells because at G2/M procentrioles are elongated and the two duplicated centrioles become well separated due to centrosome cohesion loss.

### CS isolation

The CS was isolated from RPE1 cells based on the protocol used for centrosome isolation^46^. In brief, ten 15-cm plates of *STIL^-/-^* cells arrested at S phase were allowed to swell in ice-cold hypotonic buffer (20mM K-HEPES pH7.8, 5mM K-acetate, 0.5 mM MgCl2, 0.5mM DTT). Cells were scraped off the plates and disrupted in a Dounce homogenizer (Wheaton) to release CSs. Nuclei were pelleted. The supernatant, which contains CSs, was loaded onto discontinuous sucrose gradients (40%-50%-70%) in ultra-clear centrifuge tubes and centrifuged at 120,000 × g at 4 °C for 1 h. Sucrose gradients were then collected in fractions from the bottom of centrifuge tubes. Each collected fraction was examined for the presence of CSs by pelleting CSs onto coverslips. CSs are usually peaked in fractions collected between 70% and 50% sucrose solutions.

### Antibodies

Antibodies used in this study are listed with information on source and working dilution in parentheses – anti-γ-tubulin (mouse, clone Tu30, sc-51715, Santa Cruz, 1:500 or 1:50 for UExM), anti-hSAS-6 (mouse, clone 91.390.21, sc-81431, Santa Cruz, 1:200 or 1:50 for UExM; mouse, clone G-1, sc-376836, Santa Cruz, 1:200 or 1:50 for UExM), Alexa-Fluor 488 conjugated anti-acetylated α-tubulin (mouse, sc-23950 AF488, Santa Cruz, 1:50 for UExM), Alexa-Fluor 594 conjugated anti-hSAS-6 (mouse, sc-81431 AF594, Santa Cruz, 1:50 for UExM), anti-centrin2 (mouse, clone 20H5, 04-1624, Millipore, 1:1000), anti-centrin3 (mouse, clone 1-100, H00001070-M01, Abnova, 1:200), anti-HA (mouse, clone 16B12, 901501, BioLegend, 1:500 or 1:50 for UExM), anti-ALMS1 (rabbit, A301-815A for exon8, Bethyl Laboratories, 1:2000 for immunostaining, immunoblotting and PLA or 1:1000 for UExM; rabbit, 27231-I-AP for exon10, proteintech, 1:30 for immunoprecipitation; A301-816A for exon14/15, Bethyl Laboratories, 1:1000 for UExM; goat, TA302859 for exon23, Origene, 1:100 for UExM), anti- α-tubulin (rat, clone YL1/2, MA1-80017, Invitrogen, 1:100 for UExM; mouse, clone DM1A, T6199, Sigma 1:60000 for immunoblotting), anti-acetylated α-tubulin (rabbit, 5335S, cell signaling, 1:50 for UExM; mouse, clone 6-11B-1, T7451-200UL, Sigma, 1:500 for UExM), anti-CDT1 (rabbit, clone D10F11, 8064T, Cell signaling, 1:200), anti-CEP63 (rabbit, 16268-I-AP, proteintech, 1:1000 for immunostaining and immunoblotting, 1:200-500 for UExM, 1:125 for immunoprecipitation; mouse 66996-1-Ig, protein tech, 1:50-100 for UExM; mouse, clone OTI2H2, TA809289, OriGene, 1:200 or 1:50 for UExM; mouse, clone OTI2F8, TA809290, OriGene, 1:200 for immunostaining and PLA or 1:50 for UExM), anti-CEP83 (rabbit, HPA038161, Sigma, 1:100 for UExM), anti-CEP128 (rabbit, HPA001116, Sigma, 1:500 or 1:100 for UExM), anti-CEP135 (rabbit, ab75005, Abcam, 1:50 for UExM), anti-CEP152 (rabbit, HPA039408, Sigma, 1:200 for immunostaining, 1:100-200 for UExM; rabbit, A302-480A, Bethyl Laboratories, 1:400 for UExM; mouse, clone GT1285, GTX631486, GeneTex, 1:1000 for immunostaining, immunoblotting and PLA or 1:50-1:200 for UExM), anti-CEP192 (rabbit, A302-324A, Bethyl Laboratories, 1:50 for UExM), anti- CEP295 (rabbit, ab122490, Abcam, 1:50 for UExM), anti-C-Nap1 (rabbit, Tsou and Stearns ^47^, straight use for UExM), anti-CNTROB (rabbit, 26880-1-AP, proteintech, 1:100 for UExM), anti-CP110 (rabbit, 12780-1-AP, proteintech, 1:200 or 1:100 for UExM), anti-CyclinB1 (mouse, 4135, cell signaling, 1:200), anti-BrdU (rat, MCA2060, AbD Serotec, 1:500), anti-ODF2 (rabbit, HPA001874, 1:100 for UExM), anti-PLK4 (rabbit, gift from Andrew Holland, 1:10 for UExM; mouse, clone OTI3B10, TA810562, OriGene, 1:200 or 1:50 for UExM; mouse, clone OTI2D12, TA810507, OriGene 1:200 or 1:50 for UExM), anti-PCNT (rabbit, AB4448, Abcam, 1:1000 for immunoblotting; mouse, AB28144, Abcam, 1:200 or 1:50 for UExM), anti-STIL (rabbit, A302-441A, Bethyl Laboratories, 1:10 for UExM), Secondary antibodies conjugated with Alexa-Fluor 350, 405, 488, 594, 647, 680, CF647 or HRP (Thermo Fisher, 1:500 or 1:100 for UExM; Dako P0447 and Millipore AP307P, 1:2000 for immunoblotting).

### Immunoblotting and Immunoprecipitation (IP)

Cells were completely lysed in the urea buffer (50 mM Tris pH 8.0, 2% SDS, 8 M urea, 1 mM EDTA, 5% 2-ME, 0.05% bromophenol blue) or extracted in a modified TBNS buffer (20mM Tris pH8.0, 150mM NaCl, 5mM EGTA, 1.5mM EDTA, 0.5% NP-40, cOmplete Tablets EASYpack (Cat#04693116001)). For urea lysis, the whole cell lysates (WCL) were incubated at 65 °C for 15 min and cleared by centrifugation. For cell extraction in the TBNS buffer, the whole cell extract (WCE) was centrifuged at 14000 rpm to remove the insoluble fraction before proceeding to SDS-PAGE using NuPAGE™ 3 to 8% Tris-Acetate Mini Protein Gels (Invitrogen). For IP, WCE was incubated with antibodies followed by pulldown with protein G beads (Cytiva Cat#17061801).

### Immunofluorescence

Cells were fixed in methanol at −20°C for 10 min. Slides were blocked with PBSBT (3% bovine serum albumin (w/v) with 0.1% Triton X-100 in PBS) before incubating with the indicated primary or secondary antibodies. DNA was visualized using 500μg/ml 4ʹ,6-diamidino-2-phenylindole (DAPI; Molecular Probes). Fluorescent images were acquired on microscopes (Carl Zeiss) equipped with 63x or 100× oil objectives and a camera (ORCA ER; Hamamatsu Photonics). For visualization of S-phase cells, 30 μM BrdU was added to the culture as a 30 min pulse. After staining for centrosomal antigens, cells were fixed again with -20°C methanol for 10 min and then treated with 2N HCl for 30 min at room temperature, followed by BrdU staining with anti-BrdU antibodies. Captured images were processed with Zen (Carl Zeiss), ImageJ/Fiji with the QuickFigures plugin ^48^, Adobe Illustrator, and Adobe Photoshop.

### In Situ Proximity Ligation Assay (PLA)

In situ proximity ligation assay (PLA) is a technique using oligonucleotide-conjugated antibodies to detect protein interactions within cells by generating specific DNA signals amplified via rolling circle amplification^49^. PLA reactions were done using the reagents and protocol provided by Millipore-Sigma DuoLink. ALMS1, CEP152, and CEP63 primary antibodies and their concentrations used for PLA were described above in the antibody section.

### Time-lapse Microscopy

For time-lapse experiments, cells stably expressing GFP-centrin were grown on glass-bottom dishes and imaged on a microscope (Axiovert; Carl Zeiss) configured with a 63× phase objective, motorized temperature-controlled stage, environmental chamber, and CO₂ enrichment system (Carl Zeiss). Image acquisition and processing were performed using Axiovision software (Carl Zeiss).

### Expansion Microscopy (ExM)

We followed a published ExM protocol that had been optimized for cellular ultrastructures like cilia and centrioles to prepare samples expanded to ∼4x expansion (UExM)^25^. To achieve higher expansion (∼8x), the protocol was adapted based on general principles established in high-fold expansion microscopy approaches, particularly the Magnify expansion microscopy framework^50^. This ∼8x expansion condition was further optimized at the anchoring, gelation, and denaturation steps to enable ultrastructure-preserving and homogeneous expansion. Using these approaches, samples were prepared either at ∼4x expansion (UExM) or ∼8x expansion (X8ExM) for ultrastructural imaging. In brief, cells fixed with cold methanol or 4% paraformaldehyde on coverslips were treated with the infusion solution (1.4% formaldehyde, 2% acrylamide in PBS) for 5 hours at 37°C. The gelation solution (19% sodium acrylate, 10% acrylamide, 0.1% acrylamide/bis-acrylamide (19:1), 0.5% TEMED, 0.5% APS in PBS) was then added to infused cells in a chamber on ice for 1 min, followed by 1-hour incubation at 37°C. For denaturation, hydrogels were boiled in the denaturation buffer (200mM SDS, 200mM NaCl, 50mM Tris-HCl (pH=9)) for 90 mins. Hydrogels with denatured samples were processed for the first expansion in ddH₂O. Expanded gels were washed with PBS and cut to small pieces for staining. Both primary and secondary antibody reactions were carried out for 3 hours at 37°C in a shaker, except for SAS-6 staining, which was incubated overnight. After staining, gel pieces were fully expanded again in ddH₂O at room temperature, followed by mounting onto glass-bottom dishes (D60-30-1.5-N, Cellvis) for observation.

### Ex-dSTORM analysis

To retain the size of ExM samples during dSTORM imaging, the expanded hydrogels were incubated twice in freshly prepared re-embedding gel solution (10% (w/w) acrylamide [AA], 0.15% (w/w) N,Nʹ-methylenebisacrylamide [BIS], 0.05% (w/w) TEMED, and 0.05% (w/w) APS in ddH₂O) for 25 min each at RT with gentle shaking. Polymerization was initiated by placing the entire setup in a nitrogen-filled humidified chamber and incubating at 37 °C for 1.5 h. After polymerization, re-embedded gels were washed three times in ddH₂O for 20 min each. For Ex-dSTORM imaging, the re-embedded hydrogels were immersed in an imaging buffer containing Tris-HCl, NaCl (TN) buffer at pH 8.0, 10-100 mM mercaptoethylamine (MEA) at pH 8.0, and an oxygen-scavenging system consisting of 10% glucose (G5767, Sigma-Aldrich), 0.5 mg mL⁻¹ glucose oxidase, and 40 μg mL⁻¹ catalase. Image acquisition was performed on a custom-built setup based on a commercial inverted microscope (Eclipse Ti2-E, Nikon) equipped with a focus stabilizing system. For wide-field sample illumination, laser beams from a 637 nm laser (OBIS 637 LX 140 mW, Coherent), a 561 nm laser (Jive 561 150 mW, Cobolt), a 488 nm laser (OPSL 488 LX 150 mW, Coherent), and a 405 nm laser (OBIS 405 LX 100 mW, Coherent) were focused onto the back focal plane of a 100x oil-immersion objective (NA 1.49, CFI Apo TIRF, Nikon). During Ex-dSTORM acquisition, the 637 nm and 561 nm laser lines were operated at ∼3 kW/cm² to quench the majority of fluorescence from Dy654 and CF568. A 405 nm laser beam was used to reactivate a portion of fluorophores from the dark state. The 488 nm laser line was intermittently switched on every 800 frames to enable in situ drift correction, following the procedure as previously described^51^. Emitted fluorescence was filtered through a quad-band filter (ZET405/488/561/640 mv2, Chroma) and detected using an EMCCD camera (iXon Life 888, Andor-Oxford Instruments) with a pixel size of 83 nm. For two-color imaging, the Dy654 channel was conducted first, followed by the CF568 channel using a combination of the quad-band and a short-pass filter (BSP01-633R-25, Semrock). 15,000-30,000 frames were acquired per dSTORM image at a frame rate of 50 fps. Single-molecule localizations were identified using the MetaMorph Superresolution Module (Molecular Devices) with a wavelet segmentation algorithm. Final superresolution images were cleaned with a Gaussian filter with a radius of 0.7-1 pixel. Ex-dSTORM data were used for quantitative analysis of CEP152 and CEP63 radii shown in Fig. 4E. The center of each CEP152 or CEP63 ring was first determined by circle fitting the CEP152/CEP63 dots that form the ring using ImageJ. Individual radii, measured from the center to each CEP152/CEP63 dot, were statistically analyzed and presented as histogram distributions.

### Iterative Ultrastructure Expansion Microscopy (iU-ExM)

We followed the published iU-ExM protocol^37^. After fixation, cells on coverslips were incubated in UExM infusion solution for 3 hours at 37°C, followed by gelation in the first gel solution (10% acrylamide, 19% sodium acrylate, 0.1% DHEBA, 0.25% TEMED/APS) for 15 mins on ice and 45 mins at 37°C in a humidified chamber. Hydrogels were denatured in buffer containing 200 mM SDS, 200 mM NaCl, and 50 mM Tris base (pH 6.8) for 90 min at 85°C and expanded in ddH₂O. For intermediate immunostaining, gels were shrunk in 1xPBS, stained following the UExM protocol, and re-expanded in ddH₂O. Expanded hydrogels were then embedded in a neutral gel (10% acrylamide, 0.05% DHEBA, 0.1% APS/TEMED) by three successive 10-min incubations on ice under shaking, followed by polymerization for 1 hour at 37°C. Gels were re-infused in UExM infusion solution for 3 hours at 37°C and subjected to second gelation (10% acrylamide, 19% sodium acrylate, 0.1% BIS, 0.1% TEMED/APS), followed by the same neutral gel embedding procedure. Finally, hydrogels were dissolved in 200 mM NaOH for 1 hour at room temperature under shaking, expanded in ddH₂O, and processed for post-expansion immunostaining as described above.

### Transmission Electron Microscopy

RPE1 cells grown on coverslips made of Aclar film (Electron Microscopy Science) were fixed in 2.5% glutaraldehyde and 4% paraformaldehyde in 0.1 M sodium cacodylate buffer, pH 7.0, at room temperature for 2 minutes using 250W microwave (microwave biological sample preparation system, Pelco BioWave Pro+). After three times 1-min buffer rinses using a 250W microwave, the samples were postfixed in 1% OsO₄ in the same buffer for 2 minutes using a 100W microwave at room temperature. After three times 1 min buffer rinses using a 250W microwave, samples were dehydrated in an ethanol series and propylene oxide, infiltrated with Spurr’s resin that was used for microwave enhancing. Samples were embedded in Spurr’s resin and sectioned with a Leica Reichert Ultracut S or Leica EM UC6 ultramicrotome. The ultra-thin serial sections (∼90 nm) were stained with 5% uranyl acetate/50% methanol and 0.4% lead citrate/0.1N NaOH. An FEI Tecnai G2 Spirit Twin transmission electron microscope at 80 KV was used for viewing, and the images were obtained with a Gatan Orius CCD camera.

## Data Availability

Source data are provided with this paper. All data generated are shown in the main text, figures, or supplementary materials, which are available from the authors upon request.

## Funding

This work was supported by the National Science and Technology Council, Taiwan (Grant No. 113-2628-E-002-014-) and the Ministry of Education (MOE) in Taiwan (NTU-114L900703) for TTY, and the National Institutes of Health Grant R01 GM-088253 (USA) for MBT.

## Author Contributions

M-F. B. T. conceptualized and supervised the work. T-J. B. C., W-Q. Y., and T. T. Y. provided Ex-dSTORM, 8x UExM, and 16x UExM studies. W-N. J., W-J. W., and M-F. B. T. performed EM studies. K. O., A. S., and D. I. performed other experimental studies. M-F. B. T., T. T. Y., J. L., T. S., K. O., A. S., D. I., and W-N. J. investigated and wrote the paper.

## Competing interests

None

---

**Supplemental Fig. S1.**
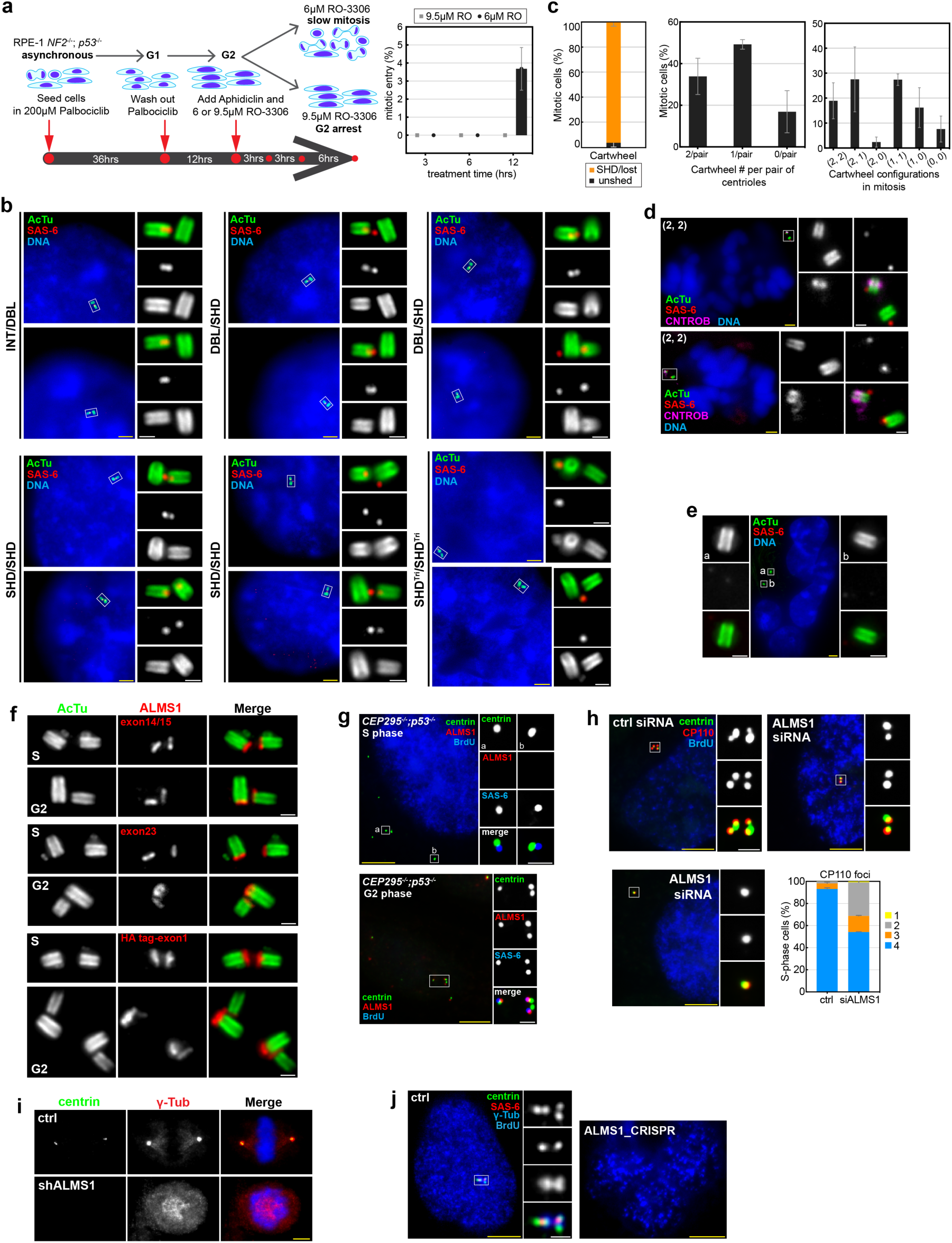
G2/M cartwheel dynamics and its association with ALMS1. (**a**) The experimental scheme of G2 cells synchronously progressing toward mitosis for a time course of 12 hrs in the presence of partial (6 μM) or strong (9.5 μM) CDK1 inhibition. Percentages of cells that had undergone mitotic entry after CDK1 inhibition at indicated time points were recorded based on time-lapse microscopy (right). While no mitotic entry was seen under strong CDK1 inhibition, ∼4% of cells had entered mitosis by the 12h timepoint under partial CDK1 inhibition. Data is mean ± s.d.. (**b**) Images of different stages of cartwheel transformation in G2 cells that slowly progressed toward mitosis under partial CDK1 inhibition. Representative images of cells hosting intact cartwheel (INT), dumbbell-like cartwheel (DBL), split-and-shed cartwheel (SHD), or shed cartwheel in centriole triplets (SHD^Tri^) were shown. (**c**) Quantification of cartwheel shedding or loss in cells that were progressing in mitosis under partial CDK1 inhibition. Left, quantification of mitotic centrioles that maintained their cartwheel in the lumen (unshed) or had shed or lost the cartwheel (SHD/lost). Middle, quantification of the cartwheel number per centriole pair in mitotic cells, ranging from 2 to 0 per pair. Right, quantification of the cartwheel configuration from two pairs of centrioles in the same cell, giving rise to 6 combinations as indicated. (**d**) UExM images of a mitotic cell showing the relative position of the shed cartwheel with the mother and daughter centrioles in the (2, 2) configuration as indicated in (**c**). CNTROB was used to mark disengaged daughter centrioles (magenta). Note that all shed cartwheels were outside the daughter lumen. (**e**) UExM images of a G1 cell that had gone through disruptive mitosis (multiple nuclei) due to partial CDK1 inhibition and had lost the intact cartwheel (no SAS-6 signals). All DNA was shown in blue (DAPI). (**f**) UExM analyses of ALMS1 localization using antibodies against ALMS1 epitopes encoded by exon14/15, exon23, or the HA epitope endogenously tagged at exon1 of ALMS1. In all cases, ALMS1 was absent at S-phase procentrioles but was localized to the proximal end of G2 procentrioles. (**g**) ALMS1 localization at newborn centrioles formed in the absence of preexisting centrioles in *p53^-/-^; CEP295^-/-^* cells. Centrin and SAS-6 labeled all newborn centrioles in S or G2, whereas ALMS1 marked only those in G2. (**h**) U2OS cells treated with siRNA for 72 hrs were examined for centriole duplication in S phase (BrdU). Failure of procentriole formation in *siALMS1* cells in S phase, as revealed by centrin and CP110 staining. Quantification is shown. Data are mean ± s.d., with N = 3 independent biological replicates. (**i**) Mitotic defects of *shALMS1* cells revealed by centrin, γ-tubulin, and DAPI staining. Note that no prominent γ-tubulin foci that normally indicate microtubule asters were seen in these mitotic-arrested cells, a phenotype not explainable by centrosome loss. (**j**) Failure of cartwheel and procentriole formation in S-phase cells (BrdU) deleted of ALMS1 by CRISPR-mediated gene targeting at exon 1. All scale bars are 1 μm (white) and 5 μm (yellow).

**Supplemental Fig. S2:**
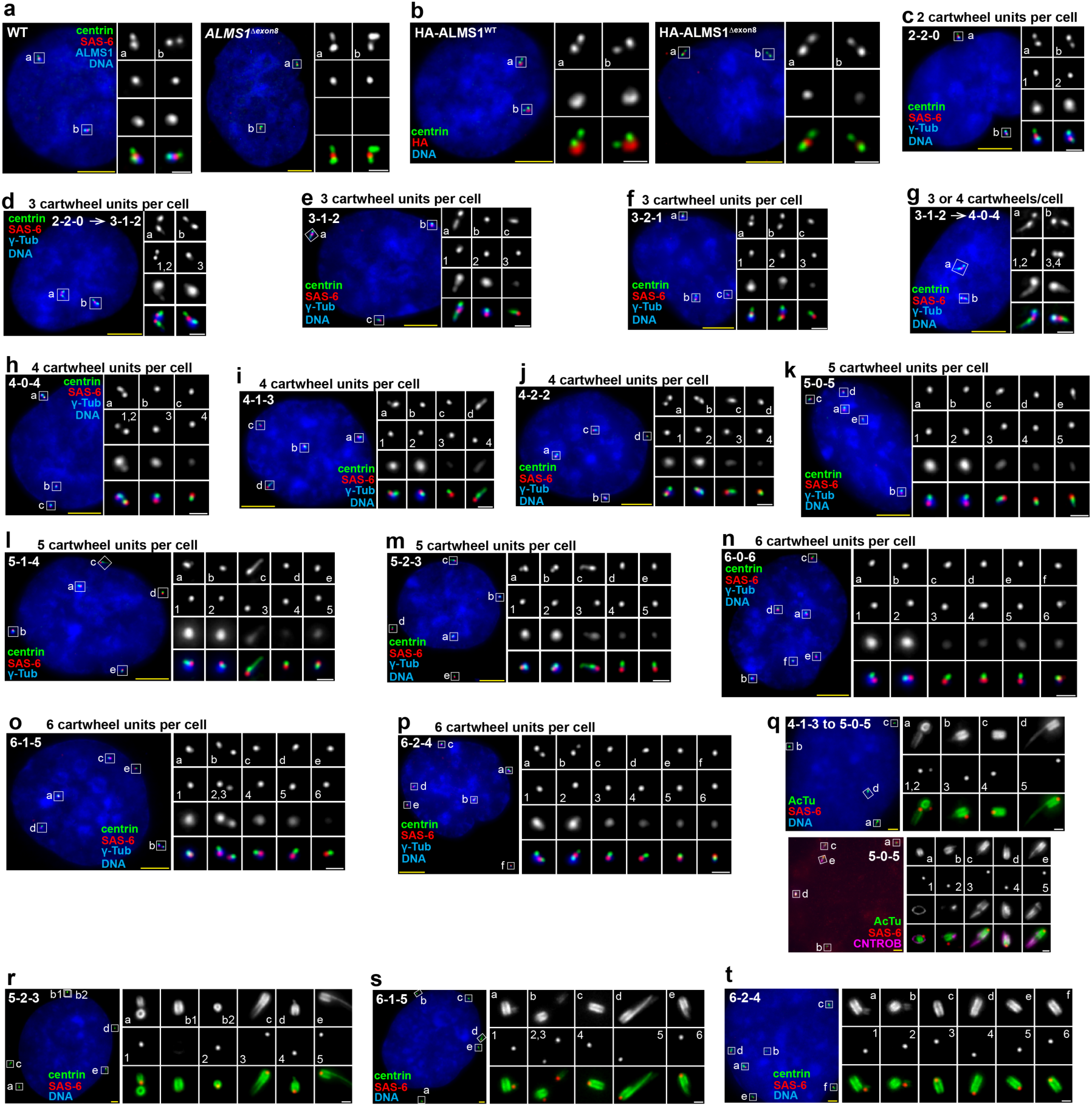
Characterization of G2/M cartwheel dynamics under *ALMS1^Δex8^* mutations. (**a**) *ALMS1^Δex8^* cells in G2 showing normal looking centrioles that had duplicated, like WT cells. Antibodies against exon8 epitopes of ALMS1 were used. (**b**) *ALMS1^Δex8^* mutant proteins, revealed by endogenously tagging HA epitopes to exon 1, were expressed and recruited to centrioles, despite at a lower level, as compared to HA-tagged ALMS1^WT^. (**c-p**) Immunofluorescence images of cartwheel amplification seen in *ALMS1^Δex8^* cells arrested in G2. The cartwheel, ranged from 2 to 6 units per cell, exhibited 12 specific configurations as illustrated in Figure 2b. Representative images revealing the transition from the 2-2-0 to 3-1-2 or from the 3-1-2 to 4-0-4 configurations were shown (**d** & **g**) (**q-t**) UExM images of 4-1-3 to 5-0-5 (**q**; top), 5-0-5 (**q**; bottom), 5-2-3 (**r**), 6-1-5 (**s**), and 6-2-4 (**t**) configurations in intact *ALMS1^Δex8^* cells. Note the difference in the relative position of the cartwheel to the mother or daughter (CNTROB); it is at the side of the mother or in the lumen of the daughter. (**q**) the 5-0-5, the cell at the top was in the transition from the 4-1-3 to 5-0-5. In the bottom cell, CNTROB labeling (magenta) marks disengaged daughters and newly formed granddaughters all carrying the cartwheel in the lumen (box c-e). (**r**) The 5-2-3: the two centrioles shown separately in b1 and b2 boxes represent a pair of engaged centrioles (a doublet) where the daughter, boxed by b2, is pointing toward the viewer. (**s**) The 6-1-5: note that cartwheel shedding and centriole disengagement occurred at one of the granddaughter centrioles, with the shed cartwheel (#2) attaching to the side of the mother, whereas the other unit (#3) remained inside the lumen of the disengaging daughter centriole. (**t**) The 6-2-4: two great-granddaughters (boxes a & b)—each growing from the shed cartwheel at the side of the mother—and 4 disengaging daughters or granddaughters (boxes c-f) each carrying a cartwheel unit in the lumen were shown. All DNA was shown in blue (DAPI). All scale bars are 1 μm (white) and 5 μm (yellow).

**Supplemental Fig. S3.**
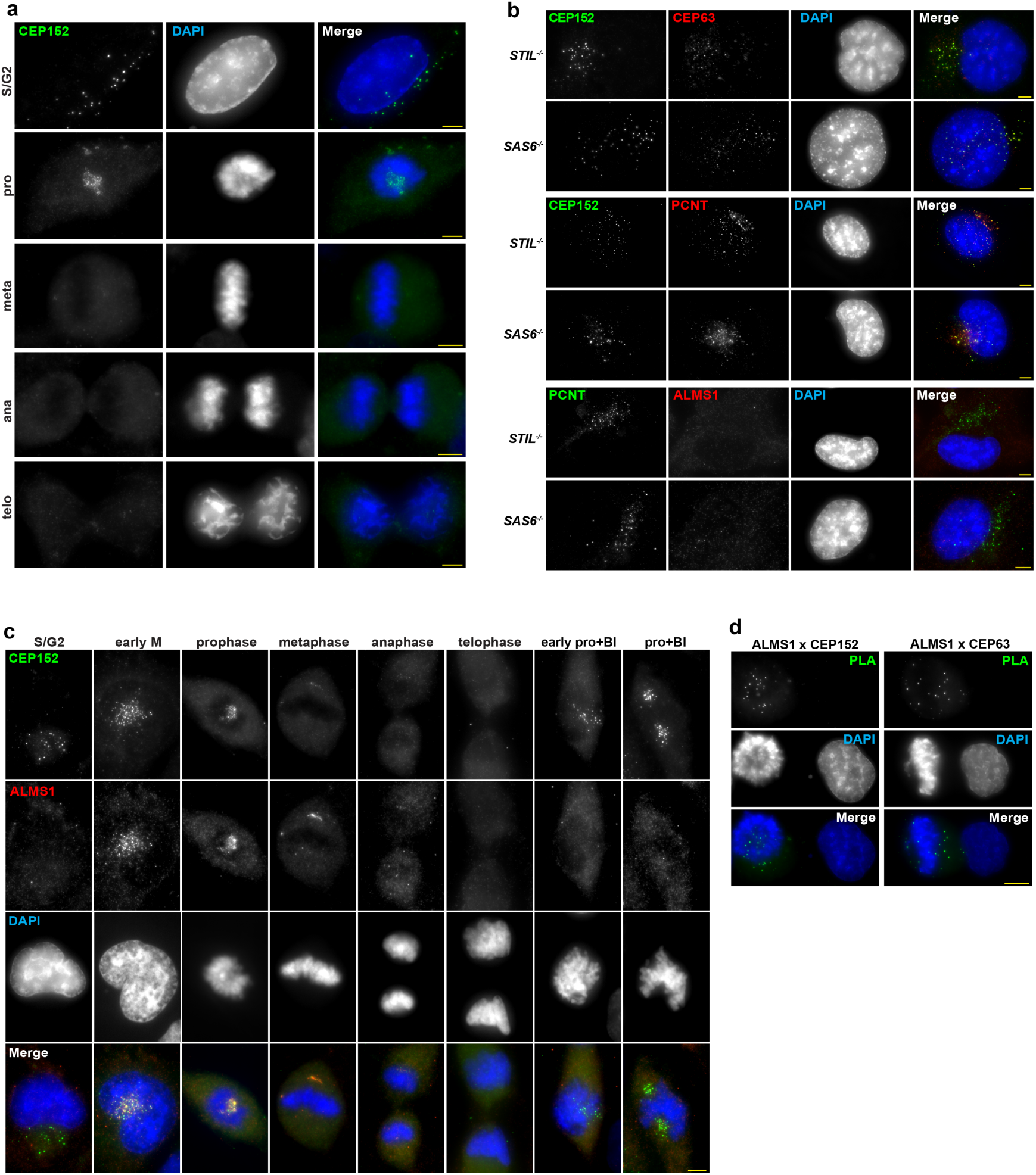
ALMS1 is physically associated with the disassembly of CEP152-CEP63-PCNT composites during the G2/M transition. (**a**) Cell cycle dynamics of CEP152 puncta shown in *SAS-6^-/-^* cells. (**b**) CEP152, CEP63, and PCNT form composites that are absent of ALMS1. Antibodies used were indicated. (**c**) Representative images of ALMS1 localization during the disassembly of CEP152 composites in mitosis. (**d**) Representative images of PLA reactions that detect complex formation of ALMS1 with CEP152 (left) or CEP63 (right) in mitotic and interphase cells present in the same field. All DNA was shown in blue (DAPI). All scale bars are 5 μm (yellow).

**Supplemental Fig. S4.**
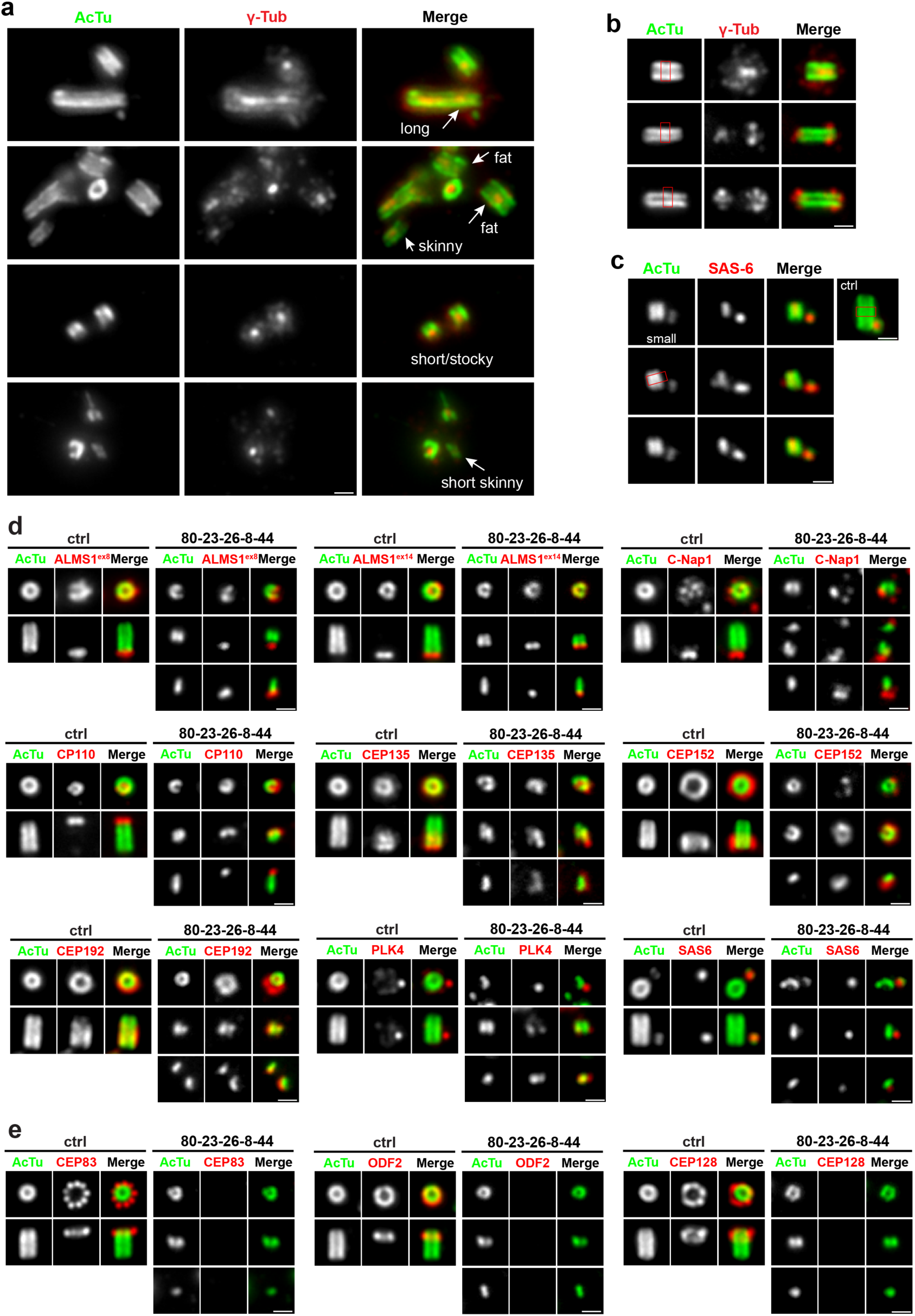
Depletion and addback of ALMS1 creates centrioles with diverse shapes that are transmittable. (**a**) Centrioles with various shapes or sizes formed after ALMS1 depletion and add-back, compared under the same magnification. (**b** and **c**) Centrioles with different shapes exhibiting distinct patterns of γ-tubulin (B) or SAS-6 (C) localization. Red boxes mark the normal width of control centrioles (**d** and **e**) Control and tiny centrioles were examined for various centriolar markers as indicated. Note that tiny centrioles cannot recruit distal or sub-distal appendages (E). All scale bars are 1 μm (white).

## Supplemental Movie

**Movie S1.**

*ALMS1^Δex8^* cells stably expressing centrin:GFP and synchronously arrested at G2 for 6 hrs were filmed by time-lapse microscopy at 5 min intervals. A G2 cell with two pairs of engaged centrioles sequentially undergoing disengagement was shown in the movie, in which centriole configuration was progressively shifted from the 2-2-0 to 4-1-3-like pattern.

**Movie S2.**

The movie shows Z-stack UExM images of CSs scattering inside a fixed *SAS-6^-/-^* cell. The CS was visualized by CEP152 (green) and CEP63 (red) labeling, with CEP152 forming ring-like structures encircling CEP63 signals. Note that due to the depth of expanded samples in gels, CSs located closer to the objective appeared brighter than those located deeper in the gel. To visualize deep CSs, surface CSs were overly intensified after image processing.

## Notes

### Competing Interest Statement

The authors have declared no competing interest.

### Summary of Updates

This revision includes new data and rewrites that clarify potential issues in the previous version.

